# Hybrid Autoencoder with Orthogonal Latent Space for Robust Population Structure Inference

**DOI:** 10.1101/2022.06.16.496401

**Authors:** Meng Yuan, Hanne Hoskens, Seppe Goovaerts, Noah Herrick, Mark D. Shriver, Susan Walsh, Peter Claes

## Abstract

**Background:** Analysis of population structure and genomic ancestry remains an important topic in human genetics and bioinformatics. Commonly used methods require high-quality genotype data to ensure accurate inference. However, in practice, laboratory artifacts and outliers are often present in the data. Moreover, existing methods are typically affected by the presence of related individuals in the dataset.

**Results:** In this work, we propose a novel hybrid method, called SAE-IBS, which combines the strengths of traditional matrix decomposition-based (e.g., principal component analysis) and more recent neural network-based (e.g., autoencoders) solutions. I.e., it yields an orthogonal latent space enhancing dimensionality selection while learning non-linear transformations. The proposed approach achieves higher accuracy than existing methods for projecting poor quality target samples (genotyping errors and missing data) onto a reference ancestry space and generates a robust ancestry space in the presence of relatedness.

**Conclusion:** We introduce a new approach and an accompanying open-source program for robust ancestry inference in the presence of missing data, genotyping errors, and relatedness. The obtained ancestry space allows for non-linear projections and exhibits orthogonality with clearly separable population groups.

## Introduction

Population structure, or the presence of systematic differences in allele frequency and linkage disequilibrium (LD) across ancestral populations, remains an important area of research in human genetics and bioinformatics. Detection thereof can be used to infer genotypic clusters, to identify admixture, and to study the history of migration and geographic isolation [1–4]. Moreover, it allows researchers to control for confounding effects due to population stratification in genome-wide association studies (GWAS), enabling accurate genetic mapping of traits [5–7].

Principal component analysis (PCA) [8] is a widely used method to analyze genome-wide single nucleotide polymorphism (SNP) data, for inferring population structure [9] and for detecting potential outliers and is routinely used as a quality control step in genetic analyses [10, 11]. In general, ancestry inference using PCA consists of the following steps. First, an ancestry space is built based on either a population diverse reference dataset with known ethnicities (e.g., HapMap project [12] or 1,000 Genome project (1KGP) [13]) solely or the combined dataset of the target population and the reference dataset. In the first approach, the target samples are projected onto the learned reference space using the same set of SNP markers. Second, the population label of an unseen target sample is inferred by applying classification algorithms, or genetic ancestry can be expressed as a continuous axis of variation. However, high-quality genotype data is critical to the success of PCA [10]. In the presence of missing genotypes or errors in the target dataset, PCA has shown to produce patterns of misalignment during projection [14, 15]. To overcome this problem, a robust alternative was recently proposed, known as SUGIBS, which utilizes spectral (S) decomposition of an unnormalized genomic (UG) relationship matrix generalized by an Identity-by-State (IBS) similarity matrix between the samples to be projected and individuals in the reference dataset [14]. By incorporating IBS information to correct for genotype errors and missing data during matrix decomposition and data projection, SUGIBS was proven to be more robust than PCA. In the work of SUGIBS, unnormalized PCA (UPCA) was investigated as well. Normalizing genotype data by allele frequency, typically done using PCA, causes individuals within the same population to be more alike, enhancing the distinction between populations [16]. This is beneficial for tasks such as clustering. However, normalization is challenging for heterogeneous datasets and increases sensitivity to outliers. UPCA (i.e., PCA on non-normalized data), on the other hand, can reduce sensitivity to outliers [14]. Another requisite of PCA is that reference subjects are unrelated to prevent fusing signals due to family relatedness with population structure [17]. For example, related individuals may form separate clusters far away from their respective ancestry groups if they were included in construction of the PCA reference space. Therefore, a common strategy is to first identify and then remove related individuals, which again requires more sophisticated solutions when dealing with heterogeneous and highly admixed datasets (e.g., KING-robust [18]).

In addition to matrix decomposition-based methods such as PCA, there has been a strong interest in the use of machine learning to identify ancestry groups and learn informative features based on genotype data. For example, support vector machines (SVM) have been applied to infer ancestry in an American population [19], and singular value decomposition (SVD) was used to reduce the dimensionality of genotype data prior to training a classification neural network to perform ancestry prediction [20]. Autoencoders (AE), a type of neural network architecture capable of learning lower-dimensional latent representations in an unsupervised manner [21, 22], have been combined with clustering methods such as K-Means and hierarchical clustering to infer population structure in maize inbred lines [23]. One advantage of neural network-based approaches is that these can easily be extended and changed. I.e., their design is flexible and today many architectural alternatives as well as training strategies exist as a result of the overwhelming scientific interest in neural network-based solutions and deep learning well beyond genetics and bioinformatics. E.g., variational autoencoders (VAE), an architectural extension of AE in terms of generative capability, were able to visualize complex population structures in a two-dimensional (2D) latent space, whereas a larger number of principal components (PCs) was required for the same task [24]. Denoising autoencoders (DAE) aim to achieve more robust latent representations by using noisy data as input and trying to reconstruct the original, clean data [25, 26]. DAE have been used for genotype imputation and provided accurate and robust results at different levels of missing data [27]. Aside from these two examples, many more alternatives exist, and this flexibility along with the ability to potentially learn non-linear relationships within the data makes neural network-based solutions very attractive to explore.

In this work, we integrate recent advances obtained for matrix decomposition-based solutions with a neural network learning paradigm into a novel hybrid approach, referred to as SAE-IBS, which consists of a Singular Autoencoder (SAE) generalized by an IBS similarity matrix (model architecture in Fig 1). This proposed method was compared with PCA and regular autoencoders through the following experiments. First, we explored the properties of the obtained low-dimensional latent representations in terms of variance-covariance structure, genetic clustering, and ancestry inference through classification. Second, we investigated the robustness of constructing an ancestry space in the presence of related individuals. Finally, we simulated missing and erroneous genotypes in the target samples and compared the robustness of different methods during the projection of target data onto a reference space. Experiments were conducted using both simulated data and real genotype data from the 1KGP [13], the Human Genome Diversity Project (HDGP) [28], and the Adolescent Brain Cognitive Development (ABCD) project [29].

**Figure 1.**
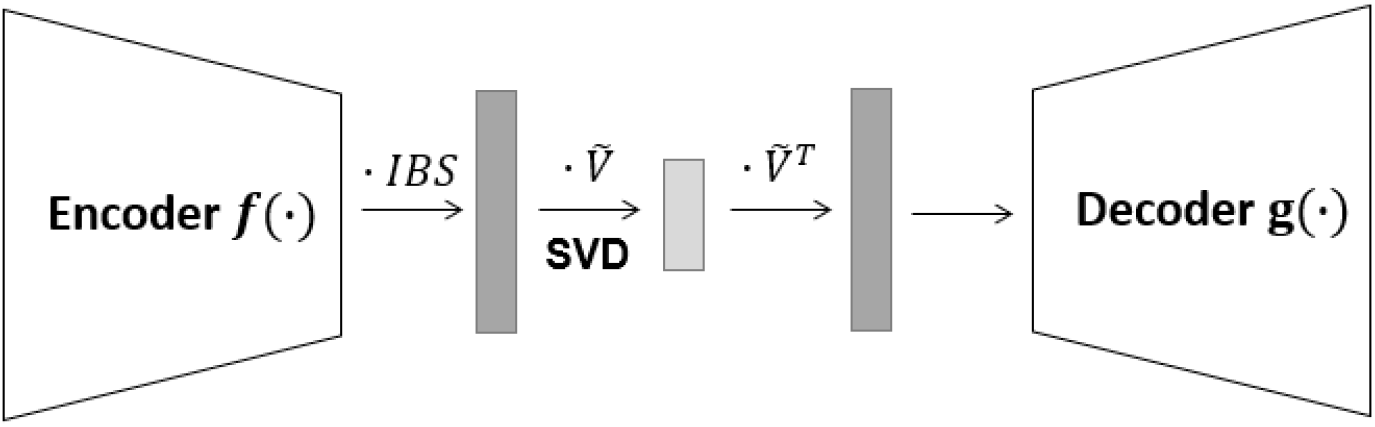
Network Architecture of SAE-IBS. The encoder and decoder are fully connected neural networks. The decoder architecture is mirrored to the encoder architecture. Matrix multiplication is denoted by · and SVD is low-rank singular value decomposition.

## Results

### Properties of the Ancestry Space

We explored the structure and properties of the ancestry spaces based on the 1KGP dataset for the three different and main categories of methods (PCA, AE, and SAE-IBS selected as representative of matrix-decomposition, neural-network, and hybrid models, respectively). More specifically, we visualized and investigated the obtained lower-dimensional latent space and its variance-covariance structure. Furthermore, we examined the discriminatory power of the latent representations for genetic clustering and classification.

Fig 2 illustrates the learned ancestry spaces of PCA, AE, and SAE-IBS. The clustering pattern resembled the geographical distribution in our data sample, with Europe, East Asia and Africa situated at different points of the triangle, South Asia positioned between Europe and East Asia, while America spread out among populations because of admixture. This structure is widely observed in ancestry spaces generated by PCA, and interestingly AE and SAE-IBS also produced similar patterns. On the other hand, samples from South Asia and America overlapped in the 2-dimensional PCA space, while training AE and SAE-IBS with only two latent axes resulted in a well-structured latent space with a clear separation of the different super-populations, which was also observed in related works such as [24, 30].

**Figure 2.**
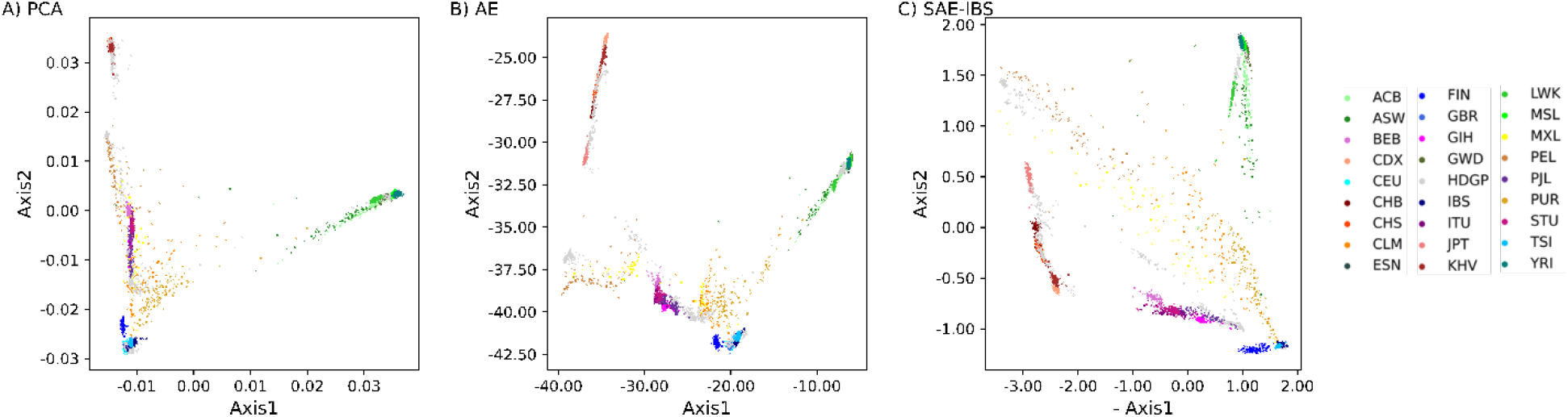
Comparison of the latent spaces for different methods. Scatter plots of the top two ancestry axes determined using (A) PCA, (B) AE, (C) SAE-IBS. The color of a point represents the ancestry of an individual, blue tints for European, green tints for African, red tints for East Asian, yellow tints for American, and purple tints for South Asian. The projected target samples from HDGP dataset onto 1KGP reference space are plotted in grey. Note in (C) SAE-IBS, to make the relative positions of clusters in different figures easier to compare visually, the first ancestry score was multiplied by -1. African Caribbean in Barbados (ACB); African ancestry in the southwestern United States (ASW); Bengali in Bangladesh (BEB); Chinese Dai in Xishuangbanna, China (CDX); Utah residents with ancestry from northern and western Europe (CEU); Chinese in Beijing (CHB); Han Chinese South (CHS); Colombian in Medellín, Colombia (CLM); Esan in Nigeria (ESN); Finnish in Finland (FIN); British from England and Scotland (GBR); Gujarati Indians in Houston (GIH); Gambian in Western Division-Mandinka (GWD); Iberian Populations in Spain (IBS); Indian Telugu in the U.K. (ITU); Japanese in Tokyo (JPT); Kinh in Ho Chi Minh City, Vietnam (KHV); Luhya in Webuye, Kenya (LWK); Mende in Sierra Leone [MSL]; Mexican ancestry in Los Angeles (MXL); Peruvians in Lima, Peru (PEL); Punjabi in Lahore, Pakistan(PJL); Puerto Rican in Puerto Rico (PUR); Sri Lankan Tamil in the UK (STU); Nigeria; Toscani in Italy (TSI); Yoruba in Ibadan (YRI).

### Variance-Covariance Structure

Fig 3 displays the variance-covariance matrix of the ancestry scores for 8-dimensional latent spaces constructed by PCA, AE, and SAE-IBS. For PCA and SAE-IBS, larger values appeared on the diagonal in descending order, indicating that PCA and SAE-IBS models capture most of the genotypic variation by the first few axes. The off-diagonal elements of the variance-covariance matrix represent the correlation between the different latent axes. For PCA and SAE-IBS, these were equal to zero due to the orthogonality that is enforced. In contrast, the latent axes of AE showed some degree of correlation, and the amount of variance was less concentrated in the first few dimensions.

**Figure 3.**
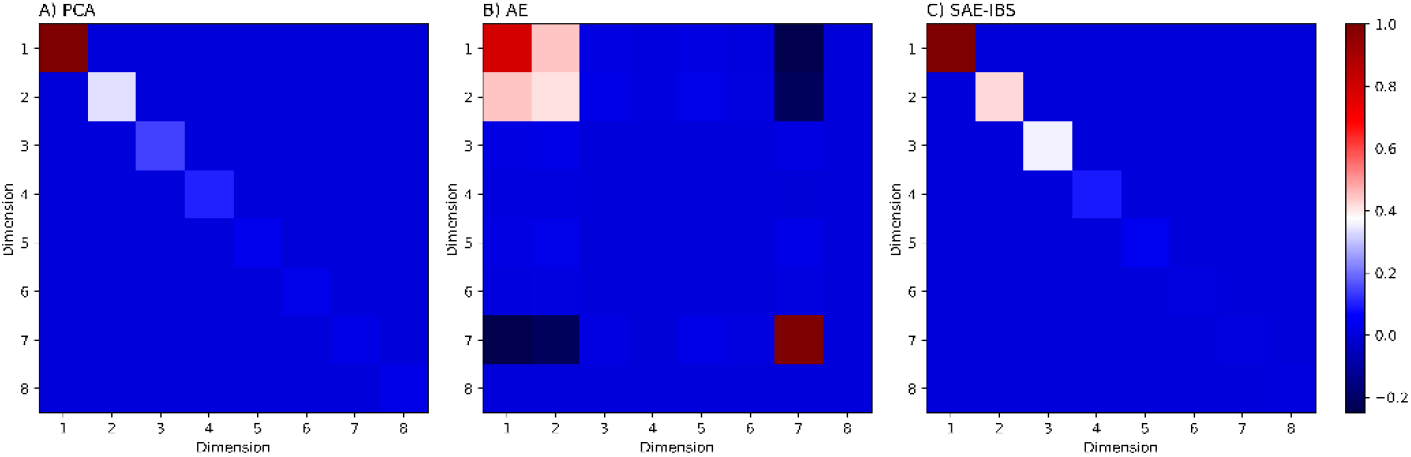
Covariance matrix of the ancestry scores for the 8-dimensional spaces constructed by (A) PCA, (B) AE, (C) SAE-IBS. The elements on the diagonal represent the amount of variance explained by each of the dimensions. The off-diagonal elements show the correlation between the axes along corresponding dimensions. Blue blocks indicate low (co)variance and red blocks indicate high (co)variance.

### Evaluation of Genetic Clustering Performance

To evaluate the validity of the different latent representations in clustering, K-means clustering [31] was applied to the 1KGP dataset with known population labels as ground truth information. Clustering performance was investigated at different resolutions, including super-population and sub-population scale. For this, the number of clusters in the K-means algorithm was set to 5 (the number of super-populations) and 26 (the number of sub-populations) respectively. Performance was evaluated using clustering accuracy (cluster labels compared to the known population labels). To investigate the influence of the number of ancestry axes on clustering accuracy, we built the models with varying latent dimensions: 2, 4, 8, and 12.

Fig 4 provides the results of clustering accuracy under different latent space dimensions of the different models. At the super-population level with latent space dimension of 2, PCA performed best, followed by the AE and SAE-IBS models. The performance of all methods increased when the latent space dimension increased from 2 to 4. However, a further increase in the number of latent dimensions reduced the performance of AE and PCA, while the performance of SAE-IBS models remained stable under different settings. The best performance of all methods was comparable and realized with 4 ancestry axes. In general, the performance for all three methods at the level of sub-population is much lower than that of the super-population. With latent space dimensions of 2 and 4, AE performed best, followed by the SAE-IBS model and PCA. The performance of PCA improved substantially when the number of latent dimensions lifted to 8, while the performance of AE and SAE-IBS models with higher latent space dimensions remained unchanged. The best performance among all approaches was realized using PCA with 8 ancestry axes. Interestingly, for PCA, the best clustering accuracy was reached with a latent space dimension set to 4 and 8 for super-population and sub-population level, respectively. This was consistent with visual inspection of the plots of subsequent PCs (Fig S1. A), in which the top four PCs captured the major continental population structure, while the subsequent PCs (PC5–PC8) accounted for additional structure in sub-populations. The latter ancestry axes of AE and SAE-IBS did not show further differentiation at the level of sub-population (Fig S1. B and C).

**Figure 4.**
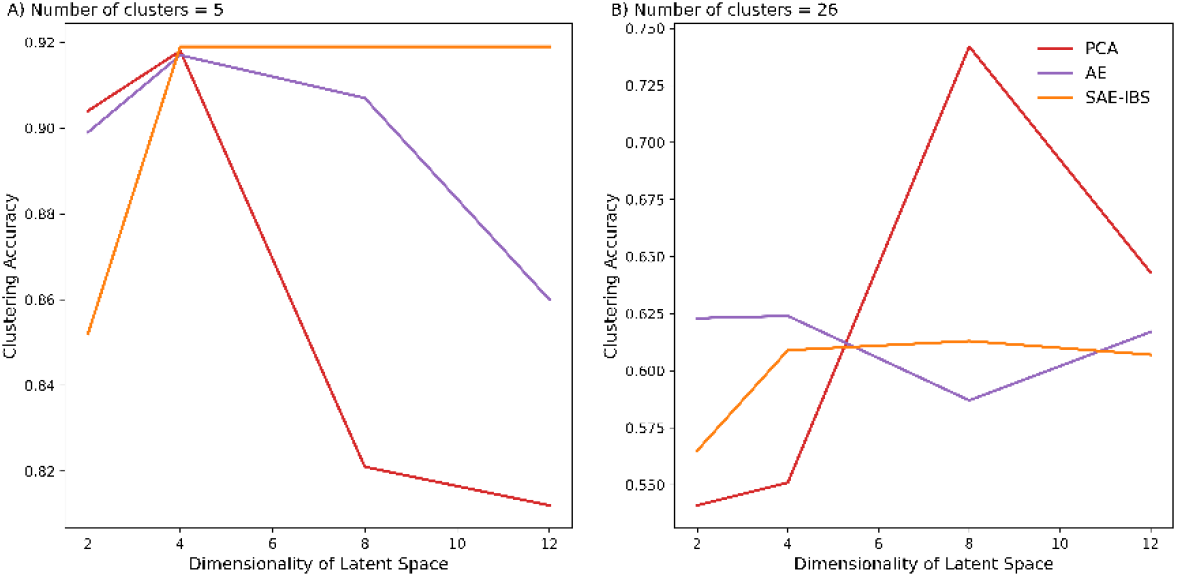
Comparison of clustering accuracy under different latent space dimensions of different models. Number of clusters in K-means algorithm was set to 5 (A) and 26 (B), corresponding to the number of super-populations and sub-populations defined in the 1KGP dataset, respectively.

Although AE and SAE-IBS did not perform as optimally as PCA for clustering a large number of closely related sub-populations, this disadvantage could be overcome by analyzing sub-populations on a smaller scale. Fig 5 displays ancestry axes for experiments using individuals from four selected 1KGP sub-populations (CHB and KHV from East Asia; GWD and YRI from Africa). The first two PCA axes defined the clusters at the super-population level while the third and fourth axes further separated the sub-populations, clearly recapitulating the results from the previous experiment. In contrast, both super-population and sub-population structures were already captured by the first two latent axes of AE, further evidenced by the eigenvalues of the correlation matrix of the latent vectors of AE. I.e., the number of eigenvalues greater than 1 (Kaiser’s rule [32]) was equal to 2, suggesting that two latent axes were adequate for clustering these four sub-populations. For SAE-IBS, the first two ancestry axes were also sufficient to reveal the hierarchical population structure, and the latter axes appeared unstructured and indistinguishable from random noise, similar to the last two axes in PCA. Furthermore, in a supplementary experiment inferencing sub-populations within one super-population (Fig S2), the first two axes of PCA, AE and SAE-IBS separated sub-populations similarly, and the last few axes of PCA and SAE-IBS appeared to capture noise. The latter axes of AE exhibited repeated structure from the first two axes, as evidenced by the variance-covariance, similar to the observations in Fig 3.

**Figure 5.**
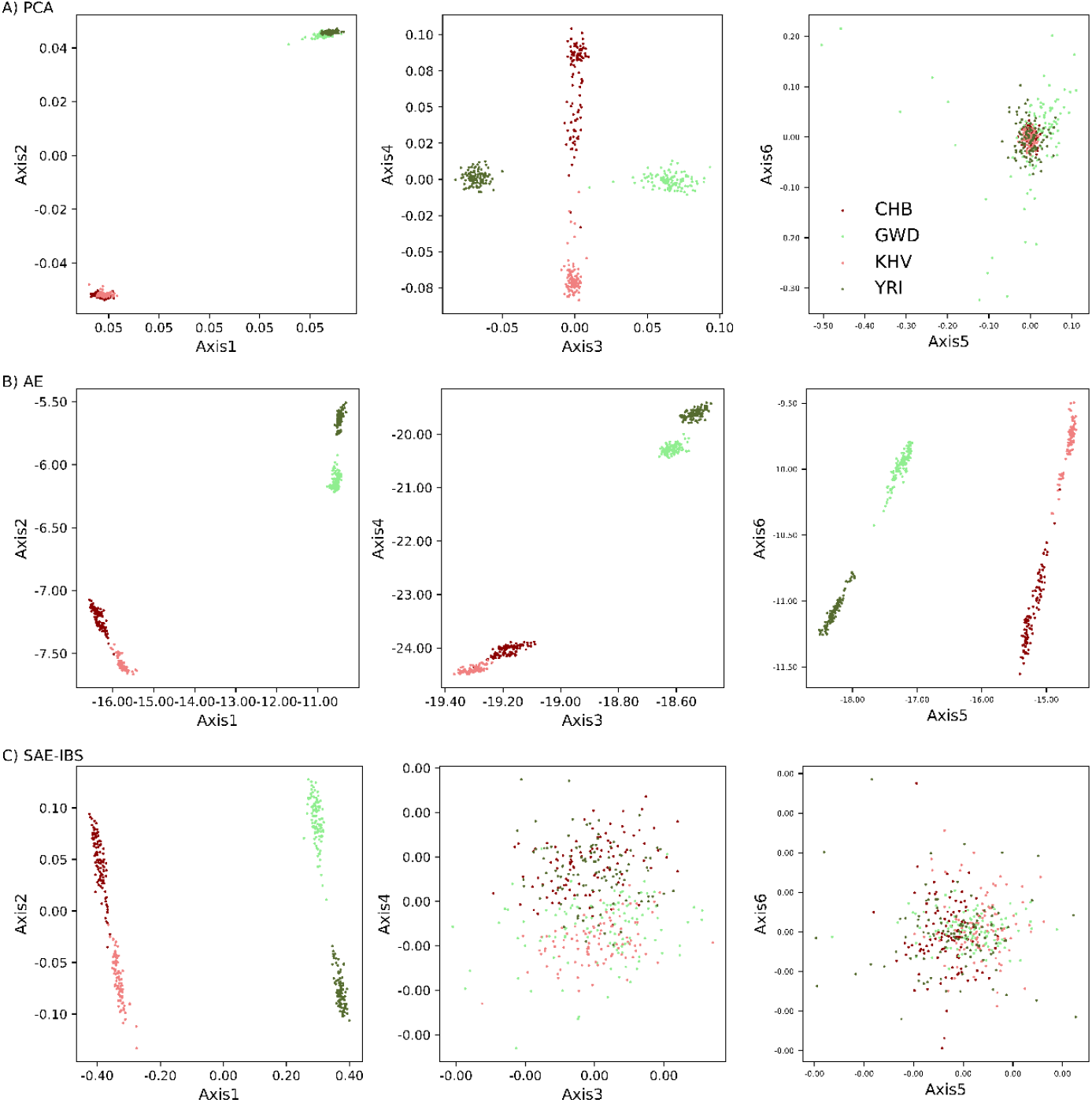
Comparison of population inference at super-population and sub-population level. The first six ancestry components for (A) PCA, (B) AE and (C) SAE-IBS using four sub-populations from the 1KGP dataset. The color of a point represents the ancestry of an individual, dark red for Han Chinese in Beijing, China (CHB), light red for Kinh in Ho Chi Minh City, Vietnam (KHV), light green for Gambian in Western Division, The Gambia (GWD), and dark green for Yoruba in Ibadan, Nigeria (YRI).

### Evaluation of Ancestry Inference Performance

The ancestry inference performance was evaluated through the classification accuracy based on the projected ancestry scores of target data. First, we built the reference ancestry space based on the 1KGP dataset. Next, a subset of the HDGP dataset was projected onto the learned space. As shown in Fig 2, the projected HDGP samples overlay well on the reference space. To further quantify the quality of these projections, we inferred the super-population labels for HDGP samples based on the labels of the 1KGP dataset using a K-nearest neighbors classification [33, 34]. Performance was assessed using classification accuracy. Since the exact match of sub-population labels between the two datasets is uncertain, we limited this experiment at the level of super-population. A summary table for sub-populations of the HDGP dataset used in this experiment can be found in Table S7.

Fig 6 provides the results of classification accuracy under different latent space dimensions for the different models. With the latent space dimension of 2, SAE-IBS performed best, followed by the AE and PCA models. The performance of all methods increased when the latent space dimension increased from 2 to 4. When the number of latent dimensions further increased, the performance of SAE-IBS remained stable, and the performance of PCA slightly improved, while the performance of AE reduced. The best performance of SAE-IBS and PCA was comparable and better than that of AE, obtained with 4 and 8 ancestry axes, respectively.

**Figure 6.**
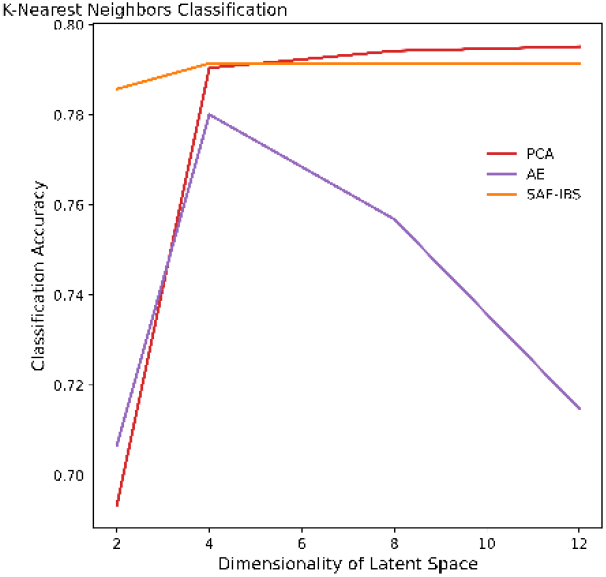
Comparison of classification accuracy under different latent space dimensions of different models.

In addition, based on the clustering and classification experiments across different dimensions of the ancestry space, we also investigated the stability of the population structure detected. By design, PCA and related matrix-decomposition solutions each time generate the same axes, independent of the dimensionality requested, with each axis being orthogonal to the others. I.e., the first four axes, when only 4 dimensions are sought for, are the same as the first four axes when up to 12 dimensions are sought for (Fig S1. A, Table S8). In contrast, the first four axes generated by an AE model trained with 4 and 12 dimensions respectively will differ (Fig S1. B, Table S8). I.e., PCA is a deterministic algorithm, while AE like many neural network methods is a non-deterministic algorithm. Interestingly, for SAE-IBS trained with 4 or 12 dimensions, the first four dimensions remain stable and are highly correlated (Fig S1. C, Table S8). This highlights a similar behavior of SAE-IBS compared with PCA, with orthogonality allowing ease of dimensionality selection for subsequent tasks.

### Construction of Robust Ancestry Space in the Presence of Relatedness

We applied the different population structure inference methods to a subset of the heterogeneous ABCD dataset consisting of unrelated individuals (n= 5,024) and 2^nd^ degree relatives detected with a relatedness threshold of 0.0884 (two different families, one of four related individuals, and one of five related individuals). With the added flexibility in training neural network solutions (AE and SAE-IBS), we investigated training an ancestry space based on the mean squared error (MSE) loss function or L2 norm (as used in the previous experiments). This makes its sensitivity to outliers equivalent to the least-square matrix decomposition encountered in PCA. In addition, we also investigated training an ancestry space based on the mean absolute error (MAE) or L1 norm. MAE represents the mean absolute distance between the original and predicted values, while the MSE measures the average of the squared difference between the original and predicted values. The squaring means the size of the error grows quadratically as the data points are located further away from the group mean. This implies that, theoretically, MSE is more sensitive to outliers than MAE.

Fig 7 displays the population structure as inferred by PCA, AE, and SAE-IBS (trained with MAE loss) with eight ancestry axes. The fifth and sixth axes of variation from the PCA model were confounded by relatedness, as shown by the related individuals that formed distinct groups far away from the main clusters. In contrast, no confounding due to relatedness was observed for AE and SAE-IBS models with MAE loss. Moreover, in a further examination with a higher latent dimension, from the 16-dimensional ancestry spaces built using AE and SAE-IBS models (Fig S3), we did not observe any separating cluster formed by the relatives. On the other hand, the obtained ancestry axes from the AE and SAE-IBS models using MSE loss were found sensitive to relatedness (Fig S4).

**Figure 7.**
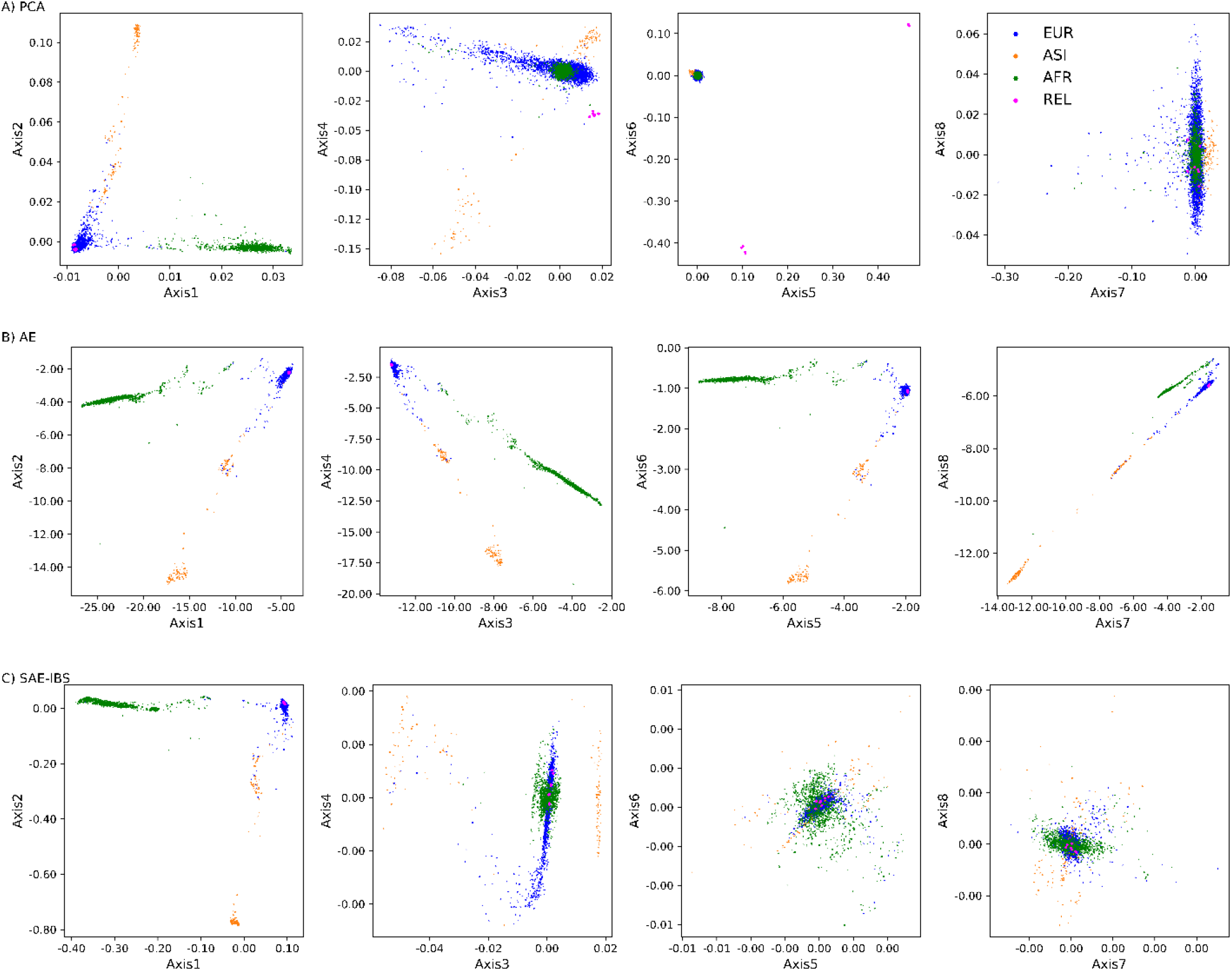
Comparison of population structure inference in the presence of related individuals. Scatter plots of the 8-dimensional ancestry space determined using (A) PCA, (B) AE, and (C) SAE-IBS, trained with MAE loss. The colors represent the self-reported ancestry of an individual, green for African (AFR), orange for Asian (ASI), and blue for European (EUR). Related individuals (REL) are plotted in pink.

To further quantify these results, we calculated the mean Mahalanobis distance (MMD)[35] between the related individuals and the three main population clusters on the first eight ancestry axes. Lower MMD values (Table 1) between the relatives and their matching ancestry group were found for AE and SAE-IBS models (trained with MAE loss) compared to PCA, indicating that neural network and hybrid solutions can be robust against related individuals within the dataset when trained with an MAE loss. Meanwhile, the robustness would be lost when the AE and SAE-IBS models were optimized in terms of MSE loss (Table S9).

**Table 1.** The mean Mahalanobis distance (MMD) using eight ancestry axes between the groups of relatives and three population clusters, i.e., European (EUR), African (AFR), and Asian (ASI). The relatives are of European descent.

| Model | MMD-EUR | MMD-AFR | MMD-ASI |
| --- | --- | --- | --- |
| PCA | 28.0230 | 151.6899 | 156.0633 |
| AE | 1.2370 | 35.7748 | 23.7082 |
| SAE-IBS | 1.8338 | 33.5951 | 30.9566 |

### Robust Projection of Target Data onto Reference Ancestry Space

To assess the accuracy and robustness against genotype missingness and erroneousness in the target data, we constructed a reference ancestry space based on the 1KGP dataset and projected unaltered and altered HDGP samples (i.e., target data of which ancestry needs to be inferred) with simulated missing and erroneous genotypes onto the reference space. Again, with the added flexibility of neural networks, in this session we also investigated denoising mechanisms for autoencoders (DAE) [25, 26] since these are relevant when working with erroneousness input data. The hybrid models (SAE-IBS, D-SAE-IBS, D-SAE-IBS-L) were compared against matrix-decomposition based (PCA, UPCA, and SUGIBS) and neural network-based (AE, DAE, and DAE-L) alternatives. DAE-L is a proposed modification of the loss function (L) in a DAE to better enforce noise robust latent representations by adding a projection loss to the reconstruction loss. D-SAE-IBS is the denoising version of SAE-IBS, and D-SAE-IBS-L is the denoising version with an extended projection loss.

Fig 8 shows the normalized root mean squared deviation (NRMSD) between the ancestry scores, computed from the original (unaltered) and simulated (altered) HGDP data, on the first two ancestry axes generated by matrix decomposition-based methods (PCA, UPCA, SUGIBS), neural network-based approaches (AE, DAE, DAE-L) and hybrid models (SAE-IBS, D-SAE-IBS, D-SAE-IBS-L). The corresponding mean and standard deviation of the NRMSD scores over 100 simulations for each experiment can be found in additional file 3 and Table S10 - S11. For the simulated erroneousness experiments, AE performed better than PCA and comparably to UPCA and SUGIBS. The denoising effect of DAE further improved the performance of AE. Moreover, DAE’s extension with an additional projection loss (DAE-L) enhanced the robustness even more. The proposed hybrid model and its extensions achieved the best performance among the above-described methods. In the scenario of simulated missingness, the denoising variants of AE (DAE and DAE-L) exhibited a similar trend in performance: by incorporating more loss terms in the objective function, the robustness increased. However, in contrast to simulated erroneousness, the group of neural network-based methods performed worse than UPCA and SUGBIS in the presence of simulated missingness. The best performance was again realized using the proposed hybrid models. Two-sample t-tests with Bonferroni correction showed significant differences in performance among most methods (p<0.0014, Table S12 - S13). For the hybrid models, however, the gain from the denoising mechanism was not substantial. I.e., SAE-IBS compared to using a denoising learning strategy in D-SAE-IBS, and further including an additional project loss in D-SAE-IBS-L did not significantly improve the results. Since the learning challenge is easier in SAE-IBS from a computational perspective, SAE-IBS may be a preferred method for this comparison.

**Figure 8.**
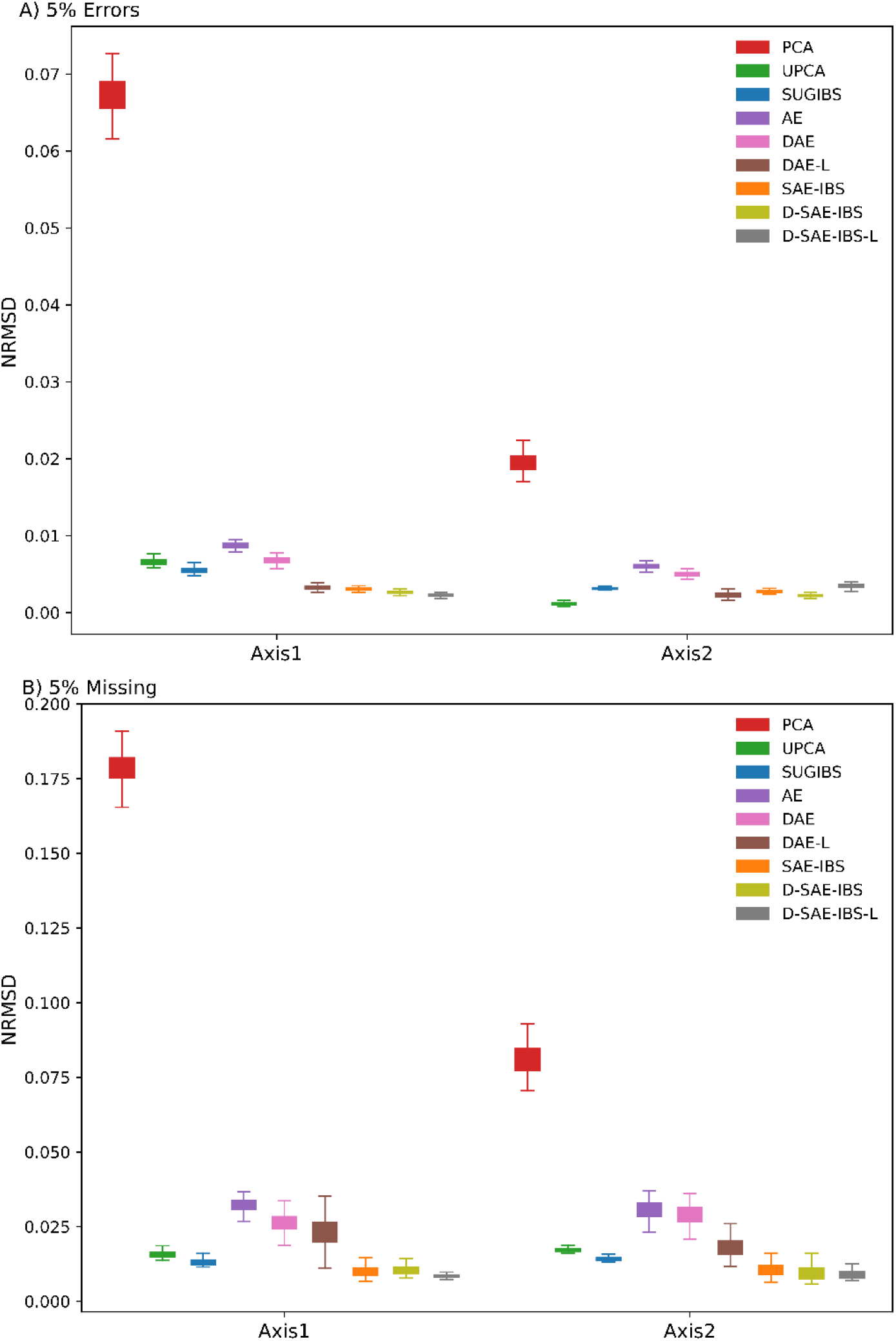
Normalized root-mean-square deviation (NRMSD) of the first two ancestry axes of different methods. (A) simulated erroneousness experiments and (B) simulated missingness experiments using matrix-decomposition methods (PCA, UPCA, SUGIBS), neural network-based methods (AE, DAE, DAE-L) and hybrid methods (SAE-IBS, D-SAE-IBS, D-SAE-IBS-L).

Furthermore, we compared the structure of the ancestry spaces in the experiments of ancestry inference and robust projection, on account of different hyperparameter settings used. As PCA generates a lower-dimensional space via matrix decomposition, one and the same ancestry space is obtained for different tasks. In contrast, the flexibility of neural network-based solutions does require a wider exploration of network implementations, known as hyperparameter tuning in machine learning. The networks can be tunned according to research interests such as optimizing the ability to robustly project new data onto the ancestry space and optimizing the clustering/classification of the data within disparate population groups coherent with the given population labels. Implementation details can be found in additional file 1 and Table S1 - S6. The latent space of AE customized for robust projection (Fig S5.A) was more contractive compared to that of ancestry inference experiments (Fig 2.B). For different tasks, the latent spaces of SAE-IBS (Fig S5.B and Fig 2.C) displayed a triangular shape, similar to what is typically observed for PCA in the context of population structure inference [36–38]. Besides, we found that the latent spaces of SAE-IBS under different hyperparameter settings were similar (characterized by a triangular shape and distinguishable clusters using the top two ancestry axes), while the AE latent spaces were more random and displayed great variability under different training conditions.

## Discussion and Conclusion

In this work, we proposed a hybrid method for population structure inference, SAE-IBS, which yields an orthogonal latent space, like matrix decomposition-based methods, and can extract more comprehensive latent features by exploiting the non-linear nature of neural network-based methods. SAE-IBS encodes genotyping data onto a lower-dimensional space and learns to reconstruct the input data from the obtained latent representations. At the same time, it inherits important robustness properties of SUGIBS by applying SVD on the IBS generalized latent representations.

The obtained ancestry space of SAE-IBS exhibits interesting patterns: it has a PCA-like variance-covariance structure; similar to AE, different population groups are separable within fewer dimensions compared to PCA. Also, this layout remained consistent under different hyperparameter settings and was independent of the dimensionality used to train the model. This leads to a strong and desired stability of the population structure inference and an enhanced dimensionality selection after training. One can train SAE-IBS with a higher dimensionality and reduce it a posteriori without the need to retrain the network. This indicates that the fine-tuning process of SAE-IBS remains simple, and hence results are less influenced by the choice of hyperparameters and/or dimensionality. On the other hand, the latent spaces of AE and its extensions varied under different combinations of hyperparameters, meaning that additional efforts are needed for fine-tuning the AE towards ancestry inference or robustness in noisy data projections. A stronger weight decay regularization effect, i.e., more contractive latent space, is beneficial for the task of robust projection, while smaller emphasis on regularization is preferred for the purpose of ancestry inference. In addition, the generation of ancestry axes is not invariant to the dimensionality requested, and selection of dimensionality is therefore cumbersome. To make sensible choices, one often needs to compare the results using different latent dimensions.

Based on the genetic clustering and ancestry inference performance, the properties of the latent representations were further explored. PCA and SAE-IBS reached similar classification accuracy and were better than AE. Regarding the resolution of genetic clustering, AE and SAE-IBS were able to differentiate the continental clusters within a smaller number of ancestry axes compared to PCA. An advantage of PCA that we observed, is that its first few components can explain the variation at the super-population level, and the subsequent components account for the variability at the sub-population level. In contrast, AE and SAE-IBS cannot detect population structure in such a hierarchical manner and are only capable of inferring sub-population structure on a smaller selection. This could be attributed to minimizing the reconstruction error defined as the element-wise mean square differences between the input and reconstructed data, which might not be a precise method to describe similarity for genotyping data. Thus, the detailed information explaining the variability within clusters is smoothed out. This argument is supported by [30], in which the LD patterns of the reconstructed and true genotype data were compared. An AE-based method was found to experience difficulty in capturing the rare SNP variations. To achieve better clustering accuracy at the sub-population level, other types of networks specifically designed for clustering can be investigated. For instance, deep embedded clustering (DEC) [39], which simultaneously learns feature representations and performs K-means clustering in the feature space, might be of interest for future research.

Moreover, in the experiment of constructing an ancestry space in the presence of relatedness, we show that more robust results can be obtained by simply changing the loss function from MSE to MAE. On the contrary, optimization in PCA works in a least-squares manner and hence the relatives tend to have a larger effect and form distinct clusters themselves. Robustness against relatedness is beneficial in that it could relieve some of the burden from preprocessing.

In addition to regular AE architectures, we also investigated denoising AE (DAE) architectures and proposed an extension by imposing robust projections through the incorporation of an additional loss (DAE-L). We found that these extensions could improve the robustness of AE against genotyping errors and missing data. Furthermore, these extensions were also applied to our proposed SAE-IBS model. Results showed that denoising extensions did not significantly improve the SAE-IBS model in terms of robustness against data artifacts. A possible explanation is that SAE-IBS already performs well by combining IBS information so that the gain from the denoising effect is marginal.

Nevertheless, with SAE-IBS and other neural network-based models, we have the flexibility of e.g., modifying the model architecture, adapting the loss function or using alternative training strategies. Related works such as [30] applied a convolutional autoencoder with residual connections, which offers an alternative model architecture different from the fully connected autoencoder used in this work. The effect of training scheme was demonstrated for ancestry inference versus robust projection of samples. This flexibility also allows these methods to be easily adapted towards other tasks such as genotype imputation (e.g., by learning to reconstruct genotypes accurately from the input with missing values via denoising mechanisms as described in [27]). Still, there is room for improvement in terms of robustness. It may be of interest for future research to explore other types of neural networks, such as denoising adversarial autoencoders [40]. These types of AE utilize an additional adversarial loss to minimize error of misclassification between actual and generated samples, which may further boost its performance.

Despite the current popularity and many advantages of neural network-based methods, hyperparameter tuning remains challenging. It is important to identify which hyperparameters to focus on in order to refine the search space. For AE models, weight decay regularization not only prevents overfitting, but also improves the robustness of projection by encouraging the feature extraction process to be locally invariant of the change of the input data. It is observed that a higher weight decay regularization results in a more robust projection. Similar ideas have been presented in the work of contractive autoencoders (CAE) [41], in which another form of penalty term for localized space contraction was proposed. CAE has been proven to outperform AE and ensure that the parameters of the encoder are less sensitive to small variations in the input. In the case of linear activations, the loss functions of CAE and weight decay regularized AE are identical and enforcing the weights of networks to be small is the only way to have a contraction [41]. We used the Leaky ReLU activation function, which comprises two linear functions and strictly speaking is not linear. Even so, weight decay regularization in our case resembles the underlying mechanism of CAE.

Although this work focuses on the robustness in population inference given erroneous input data, there are certainly other applications of neural network-based and hybrid methods worthy of future investigations. In particular, following [24], incorporating variational loss allows to better sample from the ancestry space and to generate artificial genotyping data. Another type of generative networks, generative adversarial networks (GAN), have also been applied to create artificial human genomes [42, 43]. It would thus be interesting to explore these models for genotype generation and possibly combine them with SAE models and IBS information.

In conclusion, the proposed SAE-IBS model is a new and hybrid approach that integrates the advantages of matrix-decomposition (PCA and SUGIBS) and neural network-based (AE) solutions into ancestry inference from genome-wide genotype data. By incorporating IBS information, like SUGIBS, it was shown to be more robust when projecting poor quality target samples onto a reference ancestry space. Interestingly, like PCA and in contrast to AE, the stability, and therefore repeatability, of the ancestry inference is very acceptable and due to the presence of orthogonality in the latent representation dimensionality selection can be done more readily. Furthermore, like AE and in contrast to PCA, our approach can construct a robust ancestry space in the presence of relatedness. Finally, the learned latent representations reflect various properties of the data such as cluster identities. Our method performs well for genetic clustering and ancestry inference at the super-population level and improves data visualization using a lower number of dimensions.

## Material and Methods

### Genotyping and Data Processing

The datasets used in this study are (1) the 1KGP [13] dataset, consisting of 2,504 individuals from 26 populations in Europe (EUR), Africa (AFR), East Asia (EAS), South Asia (SAS), and the Americas (AMR); (2) the HDGP [28] dataset with 1,043 individuals from 51 worldwide populations; and (3) a subset from the ABCD [29] dataset, consisting of 5,033 individuals of European, African, and Asian descent.

We followed the preprocessing steps described in [14]. In brief, for the HGDP dataset, we first converted the assembly from the NCBI36 (hg18) to the GRCh37 (hg19) using the NCBI Genome Remapping Service. For the 1KGP dataset, related individuals were detected using the KING-robust kinship estimator [18] with a relatedness threshold of 0.0442, corresponding to the 3^rd^ degree relatives, after which one individual in each group of relatives was randomly retained. For the 1KGP and HGDP datasets, individuals with more than 10% missing genotypes were removed. Pruning based on LD of SNPs is recommended for PCA [9]. We performed multiple rounds of LD pruning with a window size of 50, a moving step of 5, and *r*^2^cutoff of 0.2 until no additional SNPs were removed. We then intersected the 1KGP and HGDP datasets to extract the set of overlapping SNPs and match their alternate alleles, resulting in a final set of 155,243 SNPs for analyses.

For the ABCD dataset, the same procedures for removing missing genotypes and LD pruning were conducted, resulting in a final set of 137,482 SNPs for analyses. Individuals with missing self-reported ancestry were removed. Three ancestry groups were selected, i.e., Asia (ASI), Europe (EUR), and Africa (AFR). The related individuals in the ABCD dataset were detected using the KING-robust kinship estimator [18] with a relatedness threshold of 0.0884, corresponding to the 2^nd^ degree relatives. In total, 3,564 individuals were found to have at least one relative within the dataset. There were 1,692 pairs, 57 trios, one family comprising 4 siblings, and one family comprising 5 siblings. We retained the related individuals that had more than three relatives. The final set consisted of 5,024 unrelated individuals and 9 related individuals (European descent).

### Population Structure Inference Methods

#### Matrix Decomposition-Based Methods

##### Principal Component Analysis

In the context of genetic data, PCA aims to explain the variation in allele frequencies by finding a low-dimensional linear transformation that maximizes the projected variance. To solve the PCA problem, we performed SVD on the normalized genotype matrix. Given *n*_1_ individuals and *m* SNPs, let 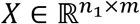 denote the unnormalized genotype matrix with additive genotype coding (aa=−1, Aa=0, AA=1 and missing=-2). The normalized genotype is obtained by 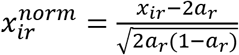, where *x*_*ir*_ and 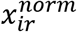 are the unnormalized genotype and normalized genotype at SNP *r* for individual *i*, respectively, and *a*_*r*_ is the sample allele frequency for SNP *r*. Then SVD takes as input the normalized genotype matrix 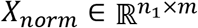 and decomposes it into a product of three matrices *X*_*norm*_ = *UΣV*^*T*^ where 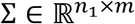 is a diagonal matrix of size *m×n*_1_ containing the singular values and the orthogonal matrices 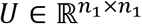 and *V* ∈ ℝ^*m*×*m*^ contain the left and right singular vectors, respectively. The dimension of the input data is then reduced by projecting it onto a space spanned by the top *k* singular vectors. Let 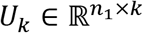 and *Σ*_*k*_∈ ℝ^*k*×*k*^ denote the left singular vectors and the singular values of the first *k*principal components, then the input data in its lower dimensional representation is given by *U*_*k*_*Σ*_*k*_, and the corresponding loading matrix is denoted by *V*_*k*_ ∈ ℝ^*m*×*k*^. The projected scores of unseen data can be obtained by multiplication of the normalized genotype matrix with *V*_*k*_.

##### Unnormalized Principal Component Analysis

UPCA works similarly to PCA, except that SVD takes the unnormalized genotype matrix as input. Interpopulation variation is captured from the second PC onwards, while the first PC represents an average SNP pattern, as is common for PCA on non-centered data. Therefore, the first PC in UPCA can be omitted.

##### Spectral Decomposition Generalized by Identity-by-State Matrix

SUGIBS was previously proposed as a robust alternative against laboratory artifacts and outliers[14] by applying SVD on the IBS generalized genotype matrix, where IBS information corrects for potential artifacts due to errors and missingness.

Let 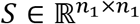 denotes the pairwise IBS similarity matrix of the unnormalized genotype matrix *X*, which is calculated following the rules in Table 2. The similarity degree of an individual is defined as 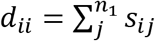 where *s*_*ij*_ is the IBS similarity between individual *i* and any other individual *j* in the reference dataset. The similarity degree matrix is a diagonal matrix defined as 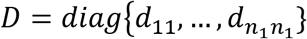. SUGIBS works similarly to PCA, except that the IBS generalized genotype matrix 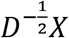 is used as input for performing SVD, i.e., 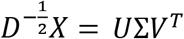. Likewise, to UPCA, the first component of SUGIBS aggregates the average SNP pattern and can therefore be omitted. For the projection of unseen samples, we use the second component and onwards 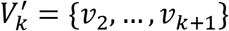 where *ν*^*k*^ is the *k*th right singular vector.

**Table 2.**
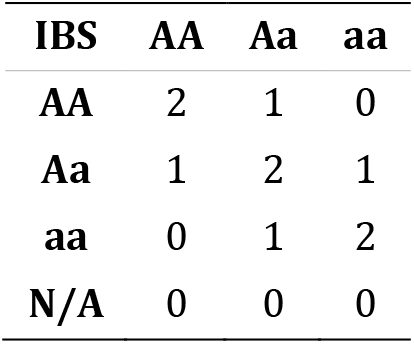
Identity-by-state similarity.

Given an unseen dataset with *n*_2_ individuals and the same set of SNPs as the reference dataset, let 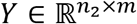 denote its unnormalized genotype matrix. The reference similarity degree is defined as 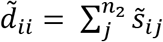 where 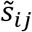 is the IBS similarity between the *i*th individual in the unseen dataset and the *j*th individual in the reference dataset. The reference similarity degree matrix is defined as 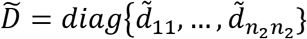. The unseen dataset can be projected onto the reference space following 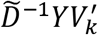.

#### Neural Network-Based Methods

##### Autoencoder

An autoencoder consists of two parts: an encoder network and a decoder network. The encoder maps input data to a latent representation *Z* = *f*(*WX* + *b*); the decoder maps the latent representation back to a reconstruction output 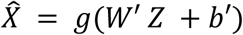 where *f*(.) and *g*(.) are nonlinear functions, *W* and *W*^′^ are the weight matrix, *b* and *b*^′^ are the bias vector, *X*, 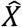 and *Z* are the input data, the reconstructed data and the latent representation, respectively. The network is then trained to minimize the reconstruction error. The objective function takes the form

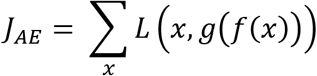

where *L* is the reconstruction error.

##### Regularized Autoencoder

To reduce overfitting of the model and improve its performance, regularization-based methods are often used. One widely used regularization is weight-decay [44], which favors small weights by optimizing the following regularized objective function

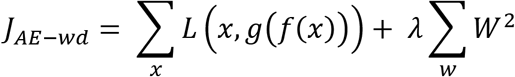

where hyperparameter *λ* controls the strength of the regularization. This encourages sparse weight matrix and thus reduces the redundancy.

##### Denoising Autoencoder

In a denoising autoencoder [25, 26], the initial input is partially corrupted before training, and then sent through the network. Based on the encoding and decoding of the corrupted input data, it is desirable to predict the original, uncorrupted data as its output. This yields the following objective function:

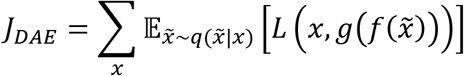

where the corrupted version 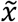 of original input *x* is obtained through the process 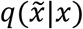.

##### Denoising Autoencoder with Modified Loss

The original objective function of a DAE focuses on denoising input, i.e., enforcing noiseless output. However, it does not guarantee learning noise-robust latent features. I.e., it is very possible that a clean and noisy input data sample are projected onto different latent representations, but still generate the same “noiseless” output. From this motivation, we propose a simple modification to the DAE’s objective function that favors robust mapping at the bottleneck/latent space. An additional term is included in the original objective function of DAE, yielding the following loss function:

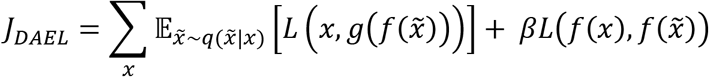

where hyperparameter *β* controls the emphasis on noise-free projections. The objective now is to learn latent representations that are not only robust for reconstruction, but also at the same time robust for projection.

#### Hybrid Methods

##### Singular Autoencoder Generalized by Identity-by-State Matrix

The general architecture of our proposed approach is displayed in Fig 1. SAE-IBS has an AE architecture, except for the bottleneck/latent space. During training, the outputs from the last layer of the encoder are multiplied by the IBS similarity degree matrix 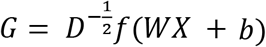 where 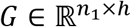 is IBS generalized features. Then, the fully connected layers linking the encoder and decoder with the bottleneck, which are typically used in regular AE, are now replaced by a low-rank SVD to ensure orthogonal latent representations. This yields 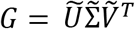 in which 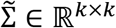 is a square diagonal matrix with its diagonal entries being the non-zero singular values of *G* and the semi-orthogonal matrices 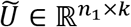 and 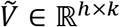 contain the left and right singular vectors, respectively. The latent representation is given by 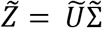, and the corresponding loading matrix is denoted by 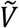. During projection, the reference IBS similarity degree matrix 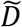 is used and the projected scores of the unseen data *Y* can be obtained by 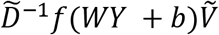. The objective function takes the form

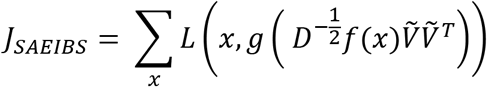

Like AE, many different variants of the general SAE-IBS architecture exist. We can extend the SAE-IBS model to a denoising version, referred to as D-SAE-IBS, by using simulated and perturbated noisy inputs and learning to reconstruct the original noise-free data input. Furthermore, we can include the extra robust projection loss in the objective function, yielding a denoising singular autoencoder generalized by Identity-by-State matrix with modified loss, referred to as D-SAE-IBS-L.

#### Implementations

##### Training Strategy

Neural network-based methods are often criticized for their danger of overfitting, i.e., the model fits extremely well to training data but fails to generalize to unseen data. To overcome this problem, we trained the models with weight-decay regularization by default. Furthermore, we applied early stopping [45] to monitor the training process and determine the optimal number of epochs. Before training, the input dataset was divided into training and validation sets (90% of samples for training, 10% for validation). After each epoch of training, we evaluated the model performance on the validation set. Training stops at the point when no improvement is observed on the validation set over a given number of epochs. The number of epochs to wait for is referred to as early stopping patience. This ensures that the training is not terminated too soon, considering the stochastic nature of training a neural network. Details of regularization and early-stopping are described in the additional file 1.

To accelerate the training process, the additive coded genotype data were first normalized so that the values are bounded within the interval of [0,1]. The encoder and decoder networks are fully connected feed-forward networks. We defined the reconstruction error as the MSE between the original genotypes and their reconstruction by default, and MAE loss was used in the experiment involving related individuals. We experimented with different model architectures, i.e., varying the number of hidden layers and the number of hidden units per layer. The model hyperparameter controlling the emphasis on projection loss *β* was also fine-tuned. The empirical analyses and detailed model architectures are described in the additional file 1.

##### Genetic Clustering

We used the K-means clustering function from the scikit-learn package. The ‘n_init’ parameter, defining the number of times to run the algorithm with different centroid seeds, was set to 1000. All other parameters were kept by default (init=‘kmeans++’, max_iter=300, tol=0.0001, verbose=0, copy_x=True, algorithm=‘auto’).

Following [46], the evaluation metric of clustering performance, i.e., clustering accuracy is defined by 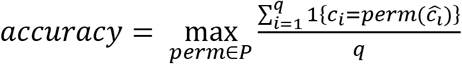 where *c*_*i*_ and 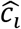 are the given true label and the predicted class label, respectively. The best assignment of the predicted labels can be found via permutation. *P* is the set of all possible permutations (in total, K! permutations) in [1, K], where K is the number of clusters. The Hungarian algorithm [47] helped to compute the best mapping between true and predicted labels efficiently, in *O*(*K*^3^).

##### Classification of Target Populations

We used the K-nearest neighbors classification function from the scikit-learn package. The number of neighbors was set to 3. All other parameters were kept by default (weights=‘uniform’, algorithm=‘auto’, leaf_size=30, p=2, metric=‘minkowski’). Classification accuracy is defined by 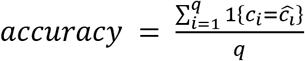 where *c*_*i*_ and 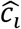 are the given true label and the predicted class label, respectively.

##### Related Individuals

In general, the top ancestry axes inferred by PCA can capture the population structure, and the latter axes might be confounded by relatedness. Therefore, the number of ancestry axes was set to 8. To quantify the distance in the multivariate ancestry space between a sample and the main population cluster, we computed the Mahalanobis distance using all 8 ancestry axes. Then, we took the mean Mahalanobis distance of all relatives as the evaluation metric of population inference accuracy.

##### Experiments of Simulated Artifacts

As shown in [14], the difference of the top eight ancestry axes across different methods exhibit similar patterns. Therefore, the number of axes to be inferred in our experiments was set to 2 for simplicity reasons only. We followed the experimental strategy described in [14], which simulates 5% genotype missingness and 5% genotyping errors (e.g., from aa to Aa or AA) of the rare SNPs (minor allele frequency < 0.05). We applied the operation of genotype masking (missingness) and alternating (errors) on the HDGP dataset to generate 100 datasets for the experiments of missingness and erroneousness, respectively. Then, the datasets with simulated noise were projected onto the 1KGP reference space. We computed the root mean square deviations (RMSD) between the top ancestry scores generated using the original (unaltered) and simulated (altered) datasets. Furthermore, to ensure comparability across methods, we normalized RMSD by the range of the scores using the original dataset, resulting in NRMSD.

To train denoising autoencoders, we generated 100 partially corrupted 1KG datasets in terms of genotype masking and alternating. In D-SEA-IBS and D-SEA-IBS-L, the IBS similarity degree matrix is calculated between the partially corrupted 1KG datasets and the target dataset. To formally test the significance of difference between different methods, we applied two-sample t-tests with Bonferroni correction on the NRMSD of the different methods over the 100 simulations. There are in total 36 independent tests, leading to a Bonferroni adjusted p-value of 0.0014 (i.e., 0.05/36).

## Abbreviations

LD: Linkage Disequilibrium
GWAS: Genome-Wide Association Study
SNP: Single Nucleotide Polymorphism
PCA: Principal Component Analysis
UPCA: Unnormalized Principal Component Analysis
SUGIBS: Spectral decomposition of an Unnormalized Genomic relationship matrix generalized by an Identity-by-State similarity matrix between unseen individuals and individuals in the reference dataset
SVM: Support Vector Machine
SVD: Singular Value Decomposition
AE: AutoEncoder
VAE: Variational AutoEncoder
DAE: Denoising AutoEncoder
DAE-L: Denoising AutoEncoder with modified Loss
SAE-IBS: Singular AutoEncoder generalized by IBS similarity matrix
D-SAE-IBS: Denoising Singular AutoEncoder generalized by IBS similarity matrix
D-SAE-IBS-L: Denoising Singular AutoEncoder generalized by IBS similarity matrix with modified Loss
1KGP: 1,000 Genomes Project
HDGP: Human Genome Diversity Project
ABCD: Adolescent Brain Cognitive Development
KNN: K-Nearest Neighbors
MSE: Mean Squared Error
MAE: Mean Absolute Error
RMSD: Root Mean Square Deviations
NRMSD: Normalized Root Mean Square Deviations
EUR: Europe; AFR: Africa; EAS: East Asia; SAS: South Asia; AMR: Americas; ASI: Asian; REL: relatives
MMD: Mahalanobis distance
CAE: Contractive AutoEncoder
CHB: Han Chinese in Beijing, China
GWD: Gambian in Western Division, The Gambia
YRI: Yoruba in Ibadan, Nigeria
KHV: Kinh in Ho Chi Minh City, Vietnam
GAN: Generative Adversarial Network
DEC: Deep Embedded Clustering

## Declarations

### Ethics statement

The datasets used in this study are openly and publicly accessible, with broad consent for research purposes.

### Consent for publication

All methods were carried out in accordance with relevant guidelines and regulations.

### Availability of data and materials

The implementation of SAE-IBS and other neural network-based methods were written in Python and available at https://github.com/mm-yuan/SAE-IBS. PCA, UPCA, SUGIBS, and simulations of erroneousness and missingness were performed using SNPLIB, a MATLAB toolbox, which can be found at https://github.com/jiarui-li/SNPLIB. The transition between human genome reference assembly was conducted via NCBI Genome Remapping Service: https://www.ncbi.nlm.nih.gov/genome/tools/remap.

The following publicly available datasets were used in this study. 1,000 Genome Project dataset https://www.internationalgenome.org/

Human Genome Diversity Project dataset: https://www.cephb.fr/hgdp/

Adolescent Brain Cognitive Development dataset: https://nda.nih.gov/abcd

### Authors’ contributions

M.Y. under supervision of P.C. and H.H. developed the SAE-IBS methodology. M.Y. together with P.C. designed the experiments with input from H.H. and S.G.. S.G. pre-processed the genotype datasets. M.Y. conducted the experiments and wrote the manuscript with extensive input from all co-authors.

## Acknowledgements

We are very grateful to Nele Nauwelaers for advice on implementing the singular autoencoder algorithm and to Dr. Jiarui Li for discussion about SUGIBS and simulation studies.

## Additional Files

### Additional file 1 — Implementation details

The encoder and decoder networks are fully connected feed-forward networks with Leaky ReLU [48] activation functions connecting each layer, except for the last layer of the decoder sigmoid activation is used to ensure the output values are bounded between [0 1]. We used the Adam optimizer [49] with an initial learning rate of 0.001. To allow the optimizer to take smaller steps when training gets close to convergence, we applied a learning rate scheduler to reduce the learning rate of the optimizer by 0.9999 after every epoch. To fit in available GPU memory (11,019MiB), we trained the networks in mini batches of 256 samples. The models are implemented and trained on an NVIDIA GeForce RTX 2080 Ti using PyTorch 1.7.

To implement the early stopping mechanism, we track if the validation loss keeps improving. If the difference of the validation loss between two epochs is below 0.1, it is quantified as no improvement. The early stopping patience was set to be 300 epochs and the maximum number of epochs equaled 3000 when training AE and SAE-IBS. For denoising extensions, every 25 epochs we generated a different simulated noisy dataset and fed to the model, therefore we relaxed the max epoch (to 5000) when training DAE, DAE-L, D-SAE-IBS, and D-SAE-IBS-L. To speed up the learning of SAE-IBS (and its denoising extensions) and to provide a well-initialized embedding from the encoder to apply SVD on, we pre-trained an AE firstly with up to 1000 epochs and continued training SAE-IBS afterward.

Following the suggestions by [50], we experimented with several parameter configurations in two steps: the first one involves the number of layers and the number of hidden units; the second one investigates emphasis on projection loss *β*. If not explicitly stated otherwise, recommended values by default in PyTorch 1.7 [51] were used for any other hyperparameters (amsgrad: False, betas: [0.9, 0.999], eps: 1e-08).

Firstly, the final hyperparameter configuration of the AE model with latent space dimension of 2 was decided. As shown in Table S1, the configuration in bold was selected as the final setting for the experiments of robust projection because it resulted in the smallest validation loss and NRMSD for the simulated missingness experiments, and relatively small NRMSD for the simulated erroneousness experiments. The same procedure was conducted for other tasks and their final settings are listed in Table S2. Then, to ensure fair comparison, the same settings were used when training AE with higher latent space dimensions, denoising variants of AE, and hybrid models. Furthermore, for the experiments of robust projection using DAE-L, we fine-tuned the hyperparameter defining the emphasis on projection loss β based on NRMSD (Table S3 and Table S4). Similarly, this parameter was tuned for D-SAEIBS-L and the final settings are displayed in Table S5 and Table S6.

**Table S1.** Comparison of different model architectures using AE with latent space dimension of 2 and weight decay of 0.01 for the experiments of robust projection. The hyperparameter configuration in bold was selected as the final setting.

| Model | Architecture | Validation loss | NRMSD missing | NRMSD error |
| --- | --- | --- | --- | --- |
| AE | 2-layer {128, 128} | 4673.00 | 0.0640 | 0.0079 |
| AE | 2-layer {512, 128} | 4668.56 | 0.0692 | 0.0091 |
| AE | 3-layer {64, 64, 64} | 4705.96 | 0.0432 | 0.0068 |
| **AE** | **3-layer {128, 128, 128}** | **4665.65** | **0.0315** | **0.0074** |
| AE | 3-layer {512, 128, 64} | 4669.04 | 0.0424 | 0.0053 |
| AE | 4-layer {128, 128, 128, 128} | 4660.43 | 0.0355 | 0.0070 |

**Table S2.**
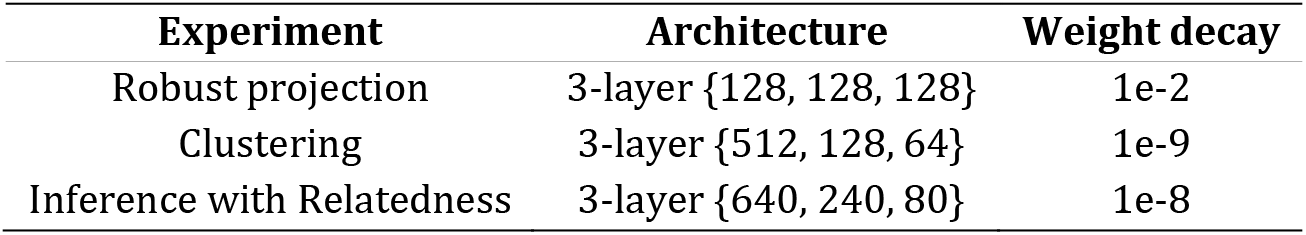
The Final hyperparameter configurations for different tasks.

| Experiment | Architecture | Weight decay |
| --- | --- | --- |
| Robust projection | 3-layer {128, 128, 128} | 1e-2 |
| Clustering | 3-layer {512, 128, 64} | 1e-9 |
| Inference with Relatedness | 3-layer {640, 240, 80} | 1e-8 |

**Table S3.** Fine-tune the hyperparameter defining the emphasis on projection loss β for the experiment of simulated erroneousness using DAE-L. The hyperparameter configuration in bold was selected as the final setting.

| Hyperparameter $\beta$ | NRMSD |
| --- | --- |
| 10 | 0.0032 |
| 50 | 0.0046 |
| 100 | 0.0052 |
| **200** | **0.0028** |
| 500 | 0.0032 |
| 1000 | 0.0040 |

**Table S4.** Fine-tune the hyperparameter defining the emphasis on projection loss β for the experiment of simulated missingness using DAE-L. The hyperparameter configuration in bold was selected as the final setting.

| Hyperparameter $\beta$ | NRMSD |
| --- | --- |
| 10 | 0.0390 |
| **50** | **0.0207** |
| 100 | 0.0276 |
| 200 | 0.0479 |
| 500 | 0.0294 |
| 1000 | 0.0366 |

**Table S5.** The Final configurations for the experiment of simulated erroneousness.

| Model | Architecture | Weight decay | Hyperparameter $\beta$ |
| --- | --- | --- | --- |
| AE | 3-layer {128, 128, 128} | 1e-2 | - |
| DAE | 3-layer {128, 128, 128} | 1e-2 | - |
| DAE-L | 3-layer {128, 128, 128} | 1e-2 | 200 |
| SAE-IBS | 3-layer {128, 128, 128} | 1e-2 | - |
| D-SAE-IBS | 3-layer {128, 128, 128} | 1e-2 | - |
| D-SAE-IBS-L | 3-layer {128, 128, 128} | 1e-2 | 100 |

**Table S6.** The Final configurations for the experiment of simulated missingness.

| Model | Architecture | Weight decay | Hyperparameter $\beta$ |
| --- | --- | --- | --- |
| AE | 3-layer {128, 128, 128} | 1e-2 | - |
| DAE | 3-layer {128, 128, 128} | 1e-2 | - |
| DAE-L | 3-layer {128, 128, 128} | 1e-2 | 50 |
| SAE-IBS | 3-layer {128, 128, 128} | 1e-2 | - |
| D-SAE-IBS | 3-layer {128, 128, 128} | 1e-2 | - |
| D-SAE-IBS-L | 3-layer {128, 128, 128} | 1e-2 | 1 |

### Additional file 2 — Additional Figures

**Figure S1.**
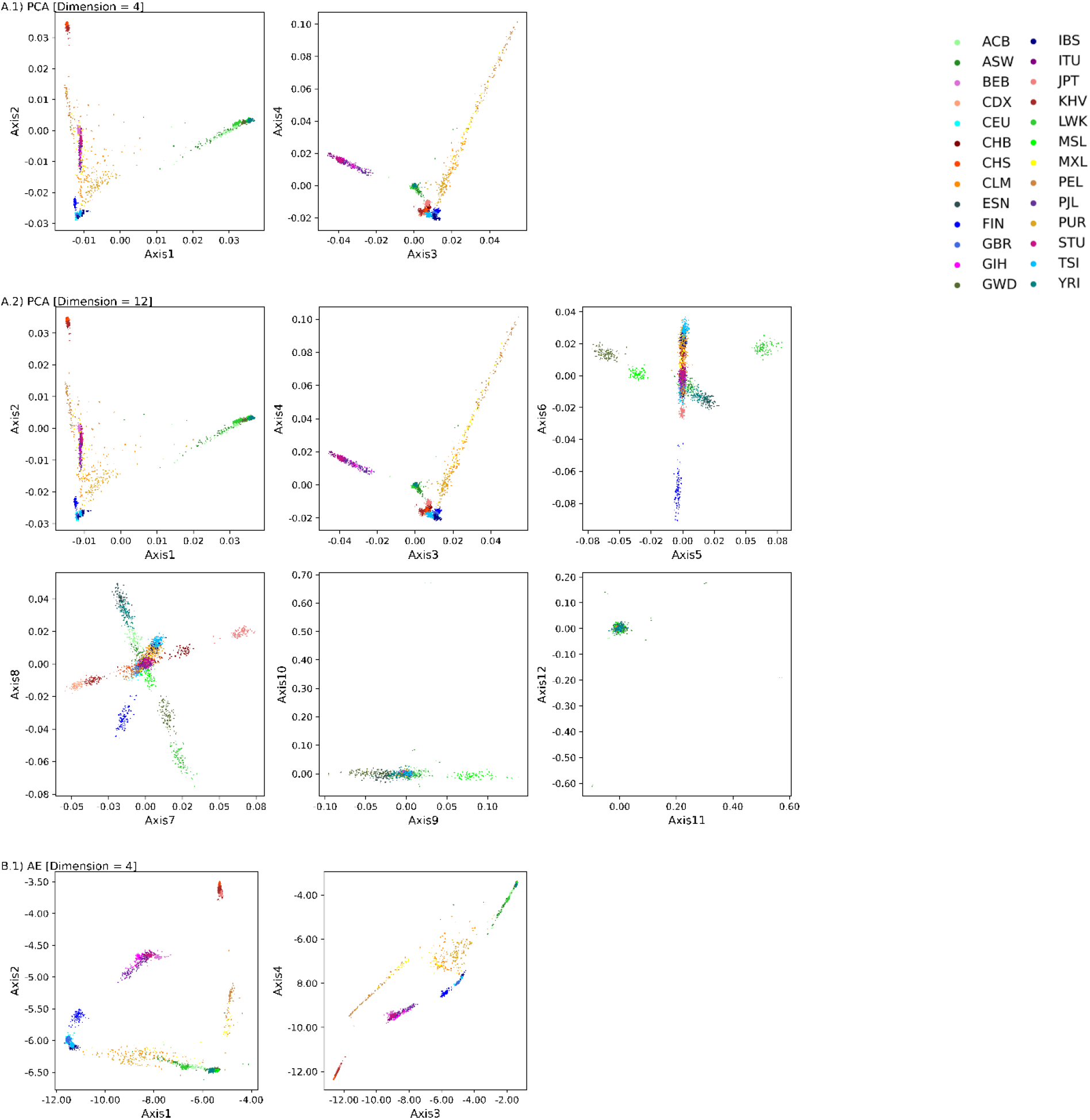
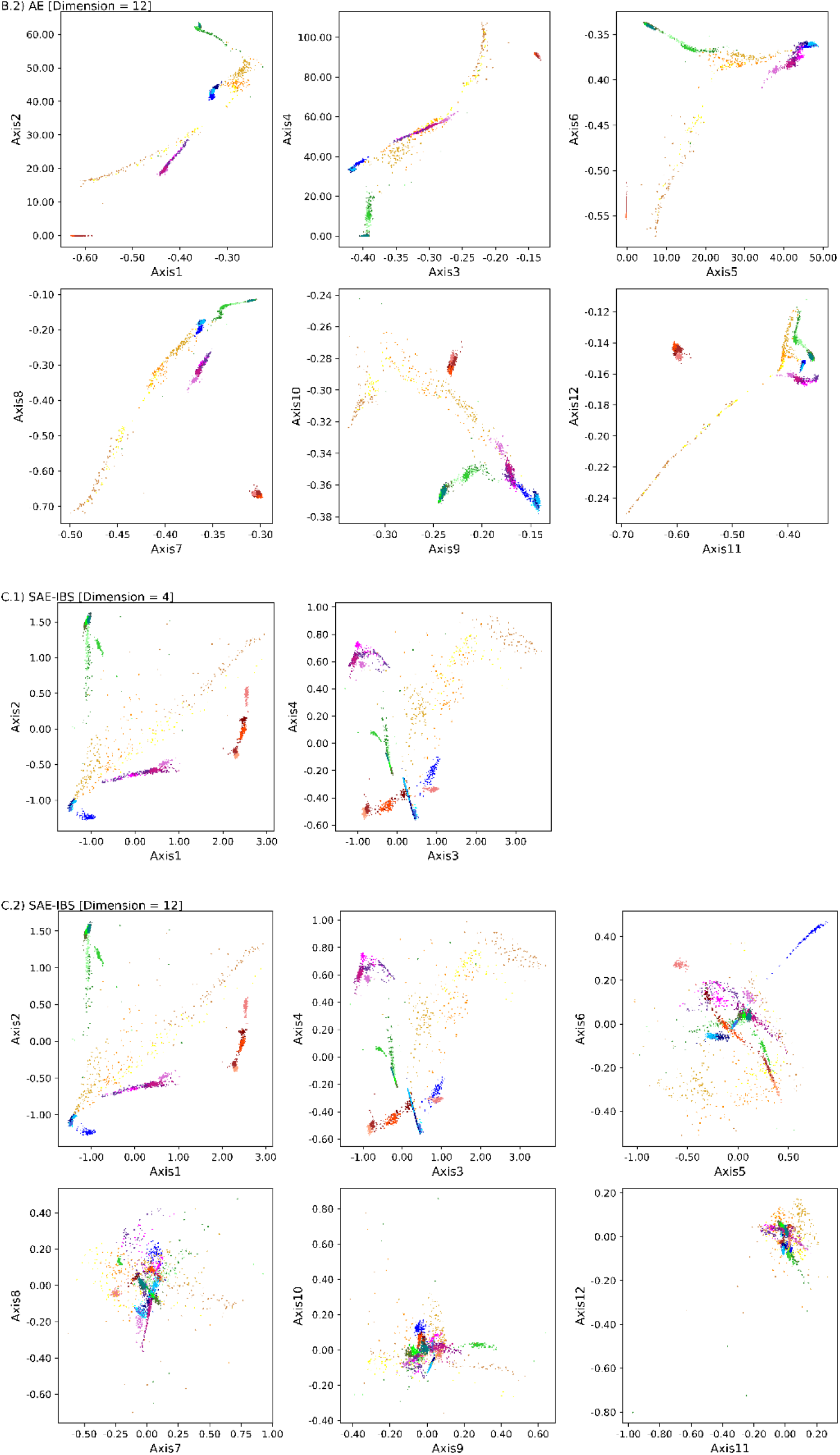
Ancestry spaces of (A) PCA, (B) AE and (C) SAE-IBS for latent space dimensions equaling 4 and 12 in the experiments of clustering and classification. The first 4 latent axes of the SAE-IBS model trained with a latent space dimension of 12 are mostly equal to the latent axes of the SAE-IBS model trained with a latent space dimension of 4. On the other hand, there is no clear pattern of latent spaces obtained using AE. The color of a point represents the ancestry of an individual, blue tints for European, green tints for African, red tints for East Asian, yellow tints for American, and purple tints for South Asian. African Caribbean in Barbados (ACB); African ancestry in the southwestern United States (ASW); Bengali in Bangladesh (BEB); Chinese Dai in Xishuangbanna, China (CDX); Utah residents with ancestry from northern and western Europe (CEU); Chinese in Beijing (CHB); Han Chinese South (CHS); Colombian in Medellín, Colombia (CLM); Esan in Nigeria (ESN); Finnish in Finland (FIN); British from England and Scotland (GBR); Gujarati Indians in Houston (GIH); Gambian in Western Division – Mandinka (GWD); Iberian Populations in Spain (IBS); Indian Telugu in the U.K. (ITU); Japanese in Tokyo (JPT); Kinh in Ho Chi Minh City, Vietnam (KHV); Luhya in Webuye, Kenya (LWK); Mende in Sierra Leone [MSL]; Mexican ancestry in Los Angeles (MXL); Peruvians in Lima, Peru (PEL); Punjabi in Lahore, Pakistan(PJL); Puerto Rican in Puerto Rico (PUR); Sri Lankan Tamil in the UK (STU); Nigeria; Toscani in Italy (TSI); Yoruba in Ibadan (YRI).

**Figure S2.**
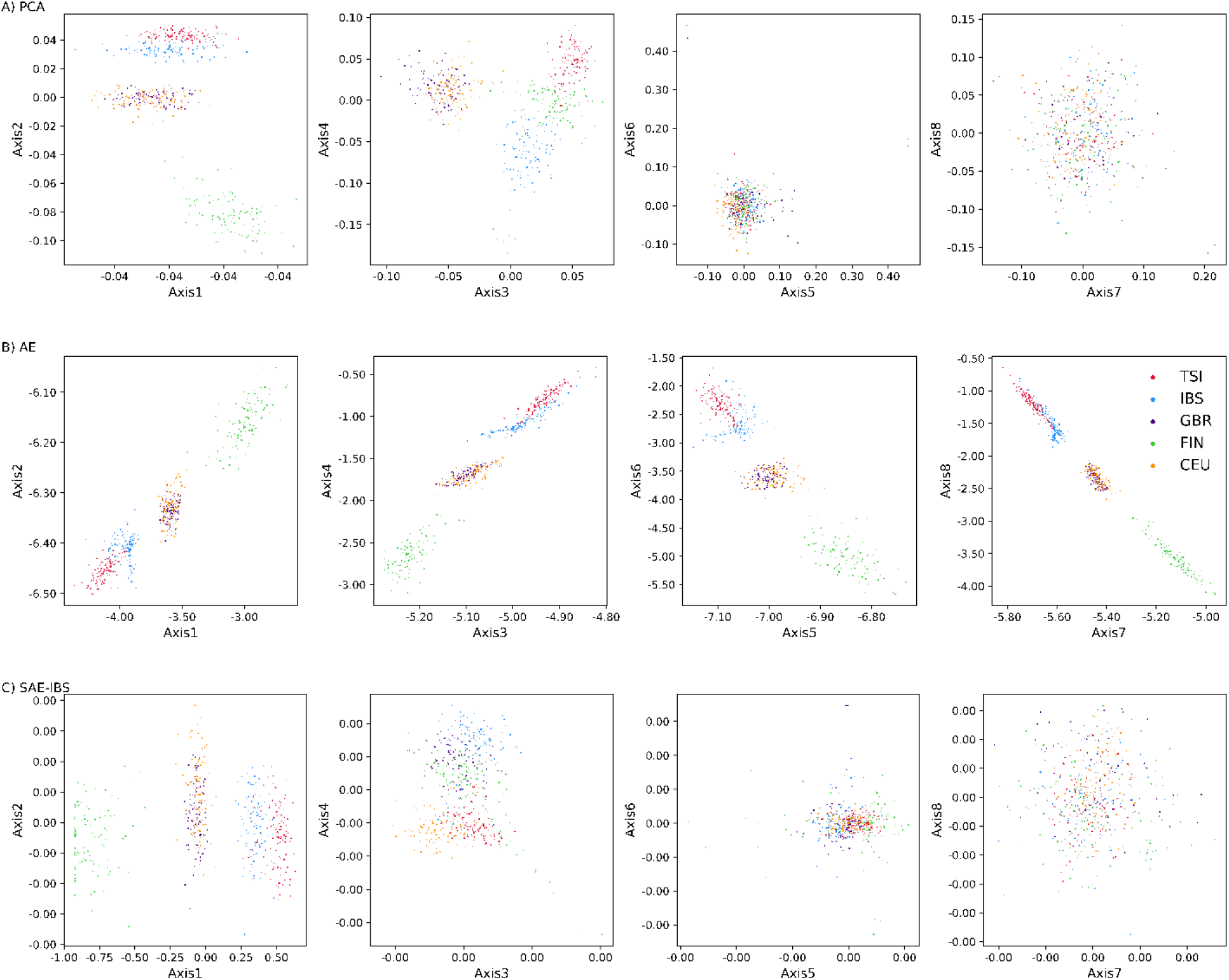
Ancestry spaces of (A) PCA, (B) AE and (C) SAE-IBS for the experiment inferencing sub-populations within one super-population. The color of a point represents the ancestry of an individual, blue for Iberian Populations in Spain (IBS), green for Finnish in Finland (FIN); red for Toscani in Italy (TSI); orange for northern and western Europe (CEU), and purple for British from England and Scotland (GBR).

**Figure S3.**
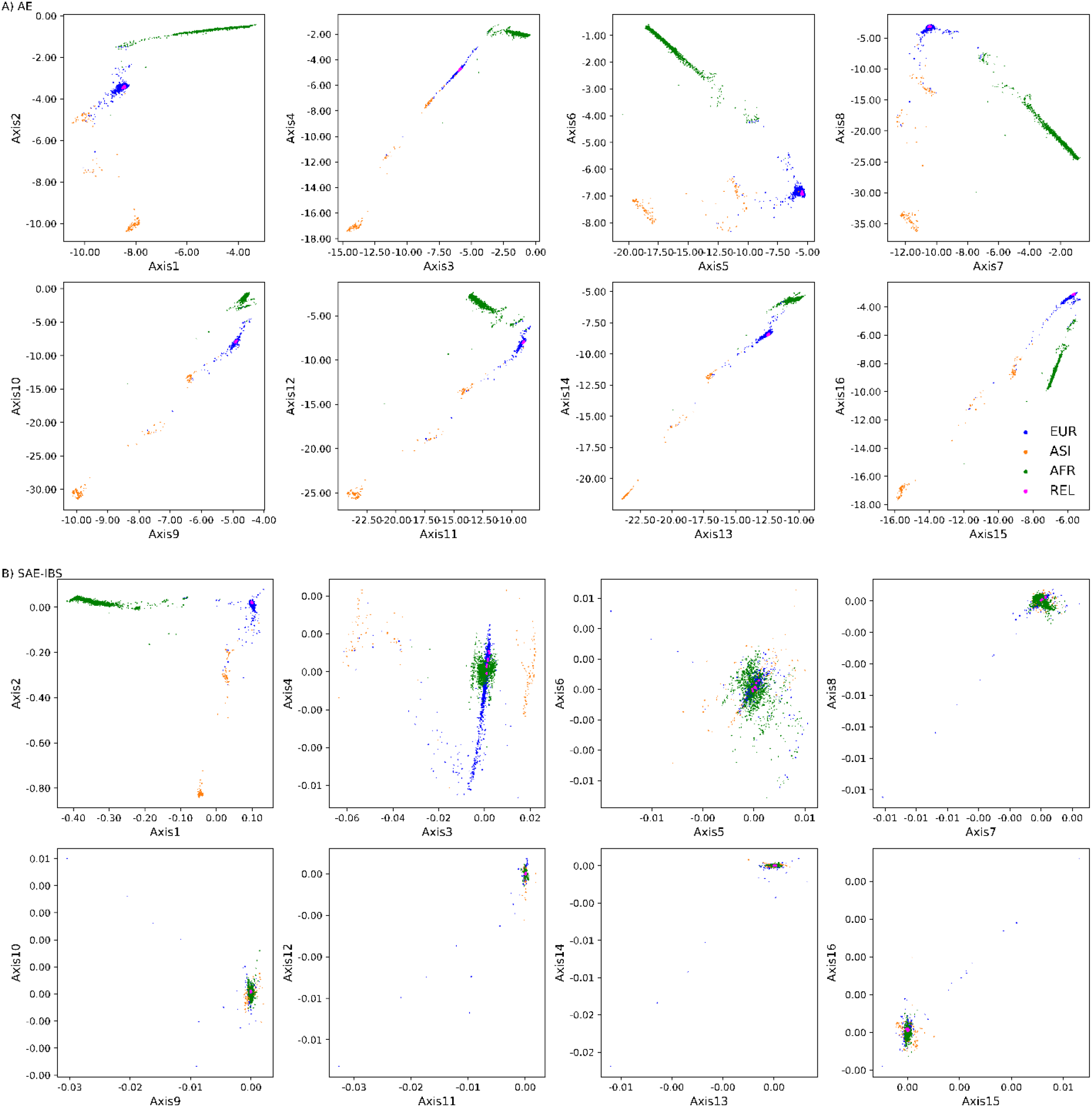
Comparison of population structure inference in the presence of related individuals. Scatter plots of the 16-dimensional ancestry space determined using (A) AE and (B) SAE-IBS, trained with MAE loss. The colors represent the self-reported ancestry of an individual, green for African (AFR), orange for Asian (ASI), and blue for European (EUR). Related individuals (REL) are plotted in pink.

**Figure S4.**
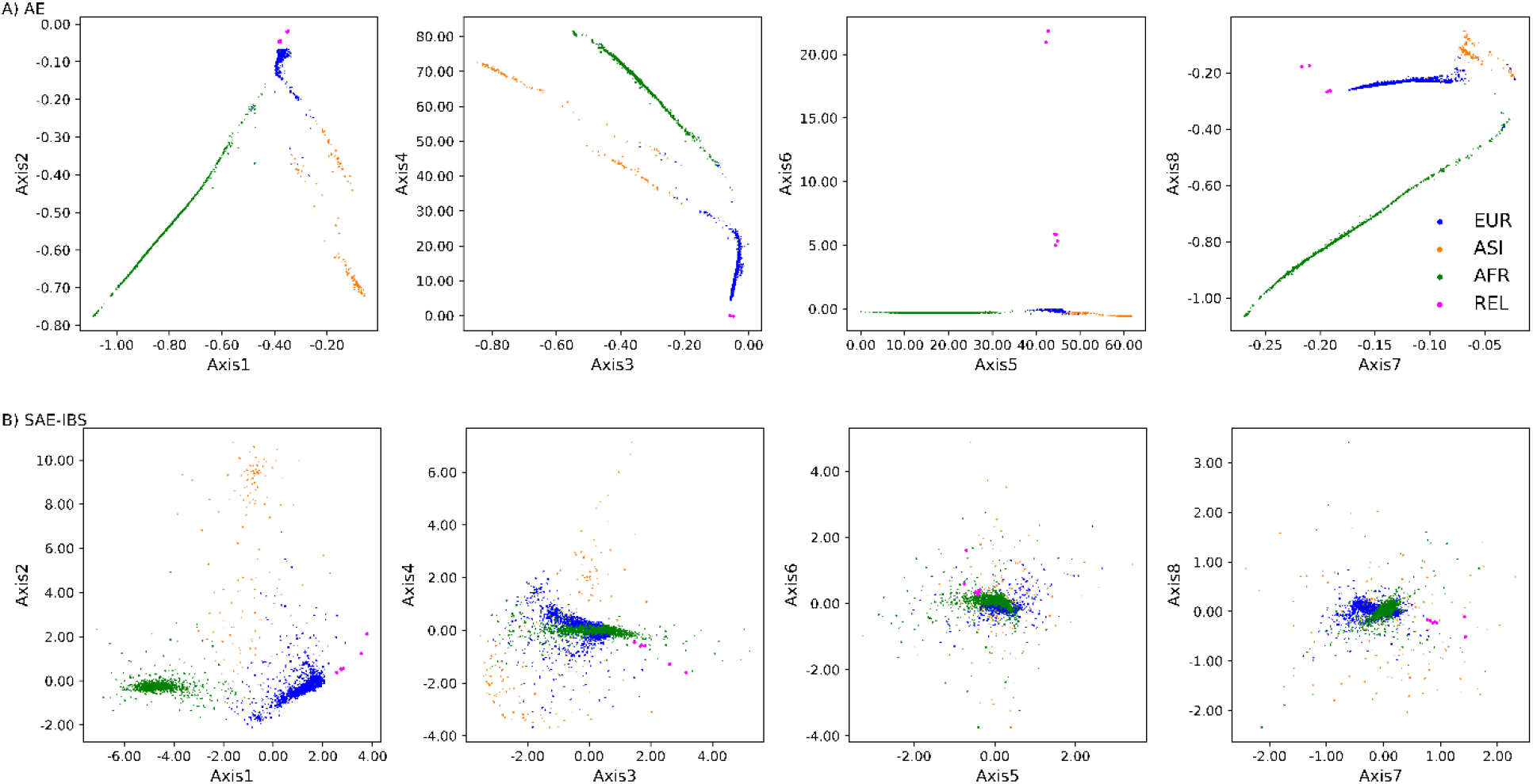
Comparison of population structure inference in the presence of related individuals. Scatter plots of the 8-dimensional ancestry space determined using (A) AE and (B) SAE-IBS, trained with MSE loss. The colors represent the self-reported ancestry of an individual, green for African (AFR), orange for Asian (ASI), and blue for European (EUR). Related individuals (REL) are plotted in pink.

**Figure S5.**
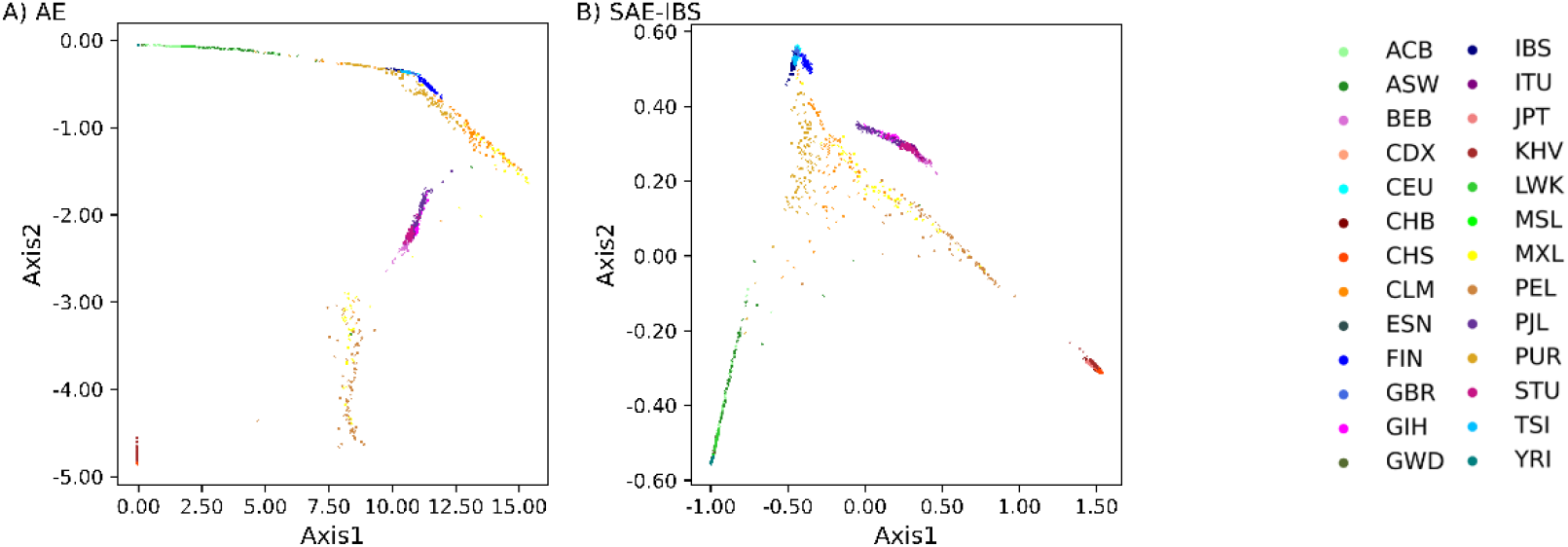
Ancestry spaces of (A) AE and (B) SAE-IBS in the experiment of robust projection.

**Figure S6.**
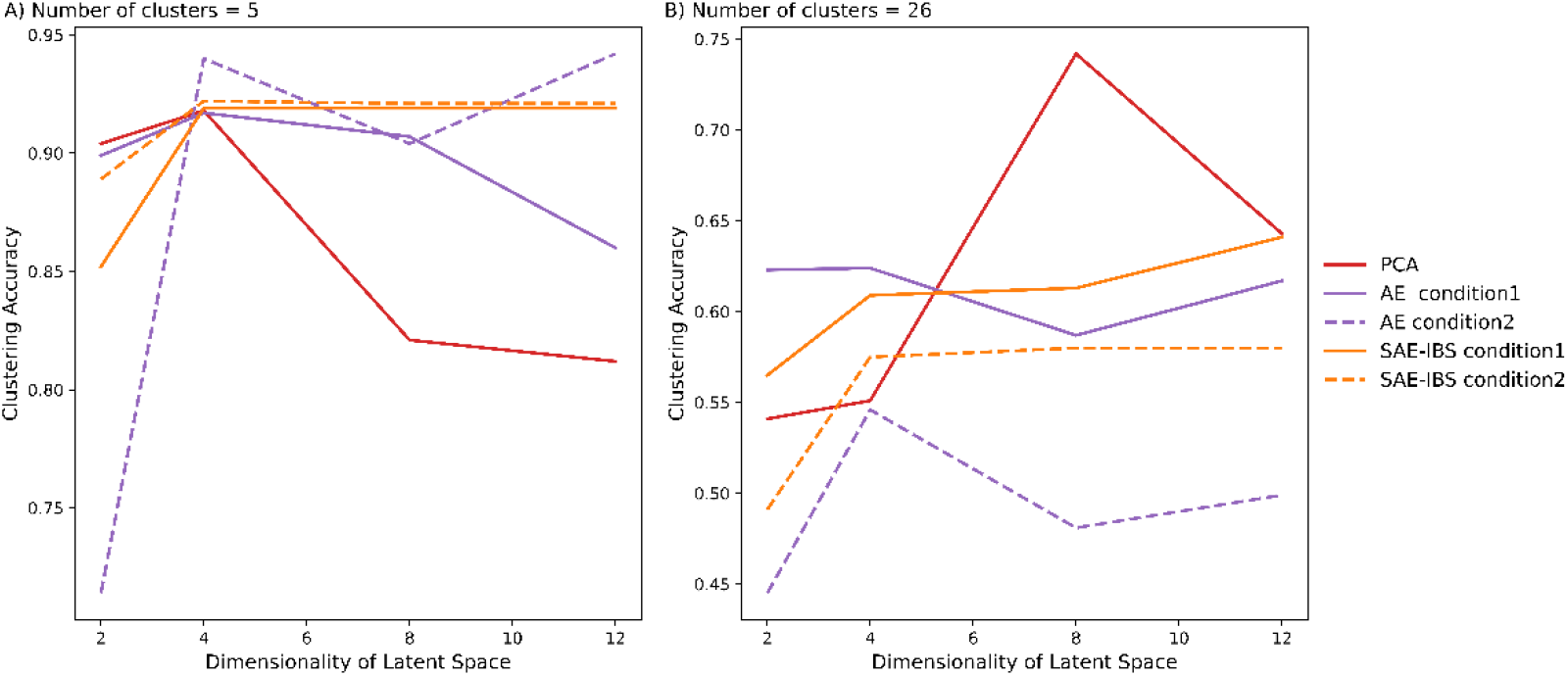
Comparison of clustering accuracy under different latent space dimensions of different models. Number of clusters in K-means algorithm was set to 5 (A) and 26 (B), corresponding to the number of super-populations and sub-populations defined in the 1KGP dataset, respectively. Condition 1 and 2 corresponds to hyperparameter settings of ancestry inference and robust projection, respectively.

### Additional file 3 — Additional Tables

**Table S7.** Summary table of the subset of HDGP dataset used in the experiment of population structure inference.

| Region | Geographic Origin | Ethnicity | Sample Count |
| --- | --- | --- | --- |
| Africa | Central African Republic | Biaka Pygmy relatives | 127 |
|  | Democratic Republic of Congo | Mbuti Pygmy relatives |  |
|  | Senegal | Mandenka relatives |  |
|  | Nigeria | Yoruba relatives |  |
|  | Namibia | San relatives |  |
|  | Kenya | Bantu NE relatives |  |
|  | S. Africa Bantu S.E. | Bantu S.E. Pedi |  |
|  | S. Africa Bantu S.E. | Bantu S.E. Sotho |  |
|  | S. Africa Bantu S.E. | Bantu S.E. Tswana |  |
|  | S. Africa Bantu S.E. | Bantu S.E. Zulu |  |
|  | S. Africa Bantu S.W. | Bantu S.W. Herero |  |
|  | S. Africa Bantu S.W. | Bantu S.W. Ovambo |  |
| South Asia | Pakistan | Brahui | 210 |
|  | Pakistan | Balochi relatives |  |
|  | Pakistan | Hazara relatives |  |
|  | Pakistan | Makrani |  |
|  | Pakistan | Sindhi relatives |  |
|  | Pakistan | Pathan |  |
|  | Pakistan | Kalash relatives |  |
|  | Pakistan | Burusho |  |
|  | China | Uygur (minority) |  |
| **East Asia** | China | Han | **216** |
|  | China | Tujia (minority) |  |
|  | China | Yizu (Yi) (minority) |  |
|  | China | Miaozu (Miao) (minority) |  |
|  | China | Oroqen (minority) relatives |  |
|  | China | Daur (minority) |  |
|  | China | Mongola (minority) |  |
|  | China | Hezhen (minority) |  |
|  | China | Xibo (minority) |  |
|  | China | Dai (minority) |  |
|  | China | Lahu (minority) relatives |  |
|  | China | She (minority) |  |
|  | China | Naxi (minority) relatives |  |
|  | China | Tu (minority) |  |
|  | Japan | Japanese |  |
|  | Cambodia | Cambodian relatives |  |
| **Europe** | France | French (various regions) relatives | **119** |
|  | France | Basque |  |
|  | Italy | Sardinian |  |
|  | Italy | from Bergamo |  |
|  | Italy | Tuscan |  |
|  | Orkney Islands | Orcadian relatives |  |
| **America** | Mexico | Pima (relatives) | **108** |
|  | Mexico | Maya (relatives) |  |
|  | Colombia | Piapoco and Curripaco relatives |  |
|  | Brazil | Karitiana (relatives) |  |
|  | Brazil | Surui (relatives) |  |

**Table S8.** Pearson correlation coefficient between ancestry axes obtained from models trained with different latent space dimensions (4 or 12). For instance, the correlation between the first axes obtained from AE trained with 4 and 12 dimensions equaled to 0.2223.

| Model | Pearson Correlation Coefficient |  |  |  |
| --- | --- | --- | --- | --- |
|  | Axis1 | Axis2 | Axis3 | Axis4 |
| PCA | 1 | 1 | 1 | 1 |
| AE | 0.2223 | 0.9251 | -0.8839 | -0.0031 |
| SAEIBS | 1 | 1 | 1 | 0.9995 |

**Table S9.** The mean Mahalanobis distance (MMD) using eight ancestry axes between the groups of relatives and three population clusters, i.e., European (EUR), African (AFR), and Asian (ASI). The relatives are of European descent.

| Model | MMD-EUR | MMD-AFR | MMD-ASI |
| --- | --- | --- | --- |
| PCA | 28.0230 | 151.6899 | 156.0633 |
| AE(MAE) | 1.2370 | 35.7748 | 23.7082 |
| AE(MSE) | 21.0173 | 5.0988E+03 | 8.8426E+03 |
| SAE-IBS(MAE) | 1.8338 | 33.5951 | 30.9566 |
| SAE-IBS(MSE) | 7.5770 | 71.9296 | 30.5254 |

**Table S10.**
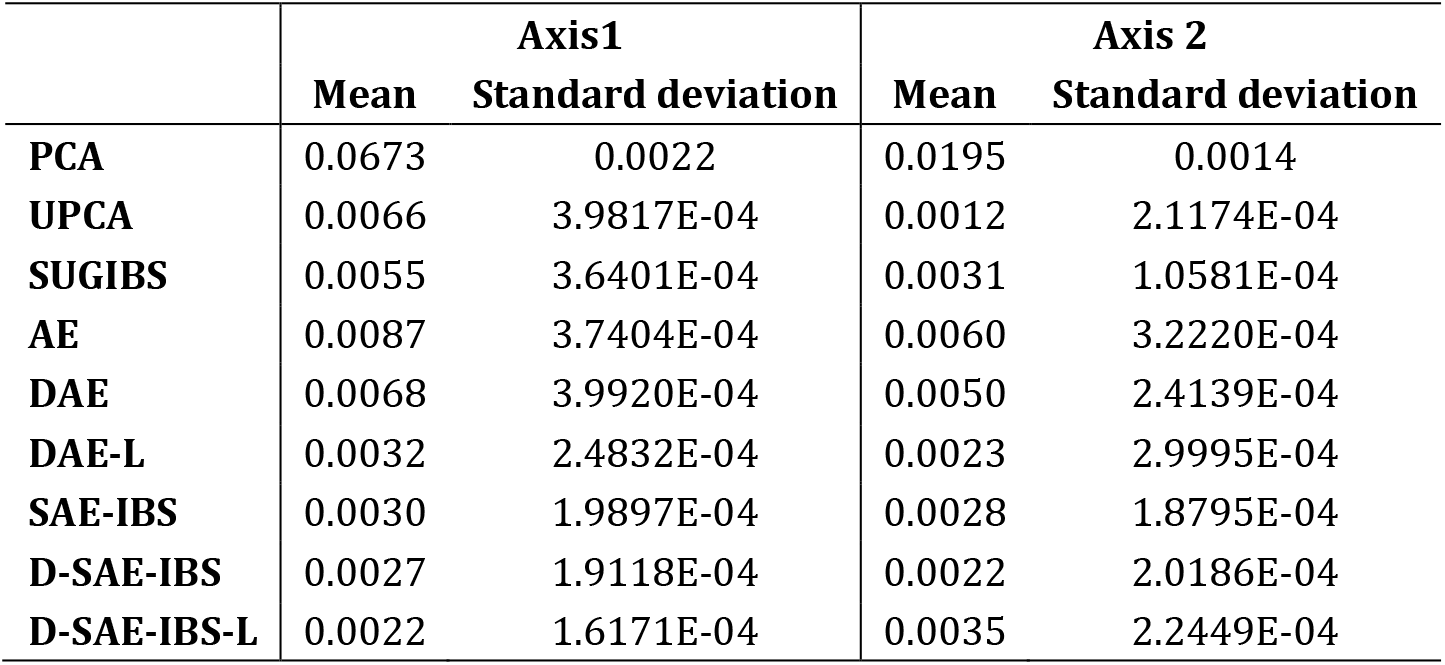
Mean and Standard deviation of the NRMSD scores over 100 simulations for the experiment of erroneousness.

|  | Axis1 |  | Axis 2 |  |
| --- | --- | --- | --- | --- |
|  | Mean | Standard deviation | Mean | Standard deviation |
| PCA | 0.0673 | 0.0022 | 0.0195 | 0.0014 |
| UPCA | 0.0066 | 3.9817E-04 | 0.0012 | 2.1174E-04 |
| SUGIBS | 0.0055 | 3.6401E-04 | 0.0031 | 1.0581E-04 |
| AE | 0.0087 | 3.7404E-04 | 0.0060 | 3.2220E-04 |
| DAE | 0.0068 | 3.9920E-04 | 0.0050 | 2.4139E-04 |
| DAE-L | 0.0032 | 2.4832E-04 | 0.0023 | 2.9995E-04 |
| SAE-IBS | 0.0030 | 1.9897E-04 | 0.0028 | 1.8795E-04 |
| D-SAE-IBS | 0.0027 | 1.9118E-04 | 0.0022 | 2.0186E-04 |
| D-SAE-IBS-L | 0.0022 | 1.6171E-04 | 0.0035 | 2.2449E-04 |

**Table S11.** Mean and Standard deviation of the NRMSD scores over 100 simulations for the experiment of missingness.

|  | Axis1 |  | Axis 2 |  |
| --- | --- | --- | --- | --- |
|  | Mean | Standard deviation | Mean | Standard deviation |
| PCA | 0.1788 | 0.0050 | 0.0811 | 0.0050 |
| UPCA | 0.0158 | 0.0014 | 0.0172 | 6.7652E-04 |
| SUGIBS | 0.0133 | 0.0013 | 0.0143 | 7.7523E-04 |
| AE | 0.0322 | 0.0023 | 0.0307 | 0.0032 |
| DAE | 0.0266 | 0.0035 | 0.0290 | 0.0035 |
| DAE-L | 0.0229 | 0.0047 | 0.0184 | 0.0038 |
| SAE-IBS | 0.0101 | 0.0019 | 0.0107 | 0.0022 |
| D-SAE-IBS | 0.0104 | 0.0016 | 0.0099 | 0.0031 |
| D-SAE-IBS-L | 0.0085 | 8.0456E-04 | 0.0093 | 0.0021 |

**Table S12.** Results of the two-sample t-tests on the NRMSD of different methods over 100 simulations for the experiment of erroneousness. Bonferroni correction method was applied to compute the adjusted significance level, accounting for multiple compassion. The non-significant p-values (p>0.0014) are marked in red.

|  | Axis1 |  | Axis 2 |  |
| --- | --- | --- | --- | --- |
|  | Mean Difference | p-value | Mean Difference | p-value |
| PCA vs. UPCA | 0.0607 | 0 | 0.0183 | 0 |
| PCA vs. SUGIBS | 0.0618 | 0 | 0.0164 | 0 |
| PCA vs. AE | 0.0585 | 0 | 0.0135 | 0 |
| PCA vs. DAE | 0.0605 | 0 | 0.0145 | 0 |
| PCA vs. DAE-L | 0.0640 | 0 | 0.0172 | 0 |
| PCA vs. SAE-IBS | 0.0642 | 0 | 0.0167 | 0 |
| PCA vs. D-SAE-IBS | 0.0646 | 0 | 0.0173 | 0 |
| PCA vs. D-SAE-IBS-L | 0.0650 | 0 | 0.0160 | 0 |
| UPCA vs. SUGIBS | 0.0011 | 1.2611E-19 | -0.0019 | 4.4342E-112 |
| UPCA vs. AE | -0.0021 | 1.2661E-66 | -0.0048 | 0 |
| UPCA vs. DAE | -2.0548E-04 | 1 | -0.0038 | 7.7335E-273 |
| UPCA vs. DAE-L | 0.0034 | 1.1542E-134 | -0.0011 | 2.1016E-44 |
| UPCA vs. SAE-IBS | 0.0036 | 3.1128E-146 | -0.0016 | 1.8354E-83 |
| UPCA vs. D-SAE-IBS | 0.0039 | 1.0473E-166 | -0.0011 | 3.7461E-41 |
| UPCA vs. D-SAE-IBS-L | 0.0043 | 6.2900E-191 | -0.0023 | 1.8843E-142 |
| SUGIBS vs. AE | -0.0032 | 1.4808E-127 | -0.0029 | 4.9692E-197 |
| SUGIBS vs. DAE | -0.0013 | 3.6642E-27 | -0.0019 | 4.0913E-110 |
| SUGIBS vs. DAE-L | 0.0023 | 3.7091E-73 | 8.2638E-04 | 4.4238E-26 |
| SUGIBS vs. SAE-IBS | 0.0025 | 4.5398E-84 | 3.3114E-04 | 2.5712E-04 |
| SUGIBS vs. D-SAE-IBS | 0.0028 | 4.2642E-104 | 8.7243E-04 | 7.5773E-29 |
| SUGIBS vs. D-SAE-IBS-L | 0.0033 | 1.0215E-128 | -3.4381E-04 | 1.1428E-04 |
| AE vs. DAE | 0.0019 | 4.4563E-56 | 9.9305E-4 | 1.3975E-36 |
| AE vs. DAE-L | 0.0055 | 5.8103E-253 | 0.0037 | 3.5246E-264 |
| AE vs. SAE-IBS | 0.0057 | 3.6417E-263 | 0.0032 | 1.0222E-224 |
| AE vs. D-SAE-IBS | 0.0061 | 6.7913E-281 | 0.0038 | 1.0051E-267 |
| AE vs. D-SAE-IBS-L | 0.0065 | 1.7604E-301 | 0.0026 | 2.3286E-167 |
| DAE vs. DAE-L | 0.0036 | 1.6086E-146 | 0.0027 | 9.4960E-183 |
| DAE vs. SAE-IBS | 0.0038 | 4.7548E-158 | 0.0022 | 2.3775E-139 |
| DAE vs. D-SAE-IBS | 0.0041 | 2.2379E-178 | 0.0028 | 1.0246E-186 |
| DAE vs. D-SAE-IBS-L | 0.0045 | 2.4796E-202 | 0.0016 | 1.6362E-80 |
| DAE-L vs. SAE-IBS | 2.0050E-04 | 1 | -4.9524E-04 | 9.3120E-10 |
| DAE-L vs. D-SAE-IBS | 5.5732E-04 | 3.0572E-09 | 4.6053E-05 | 1 |
| DAE-L vs. D-SAE-IBS-L | 9.8614 E-04 | 3.1105E-16 | -0.0012 | 4.5807E-49 |
| SAE-IBS vs. D-SAE-IBS | 3.5681 E-04 | 0.0558 | 5.4130E-4 | 1.2833E-11 |
| SAE-IBS vs. D-SAE-IBS-L | 7.8564 E-04 | 1.9353E-10 | -6.7495E-04 | 8.5458E-18 |
| D-SAE-IBS vs. D-SAE-IBS-L | 4.2883 E-04 | 0.0052 | -0.0012 | 1.6841E-52 |

**Table S13.**
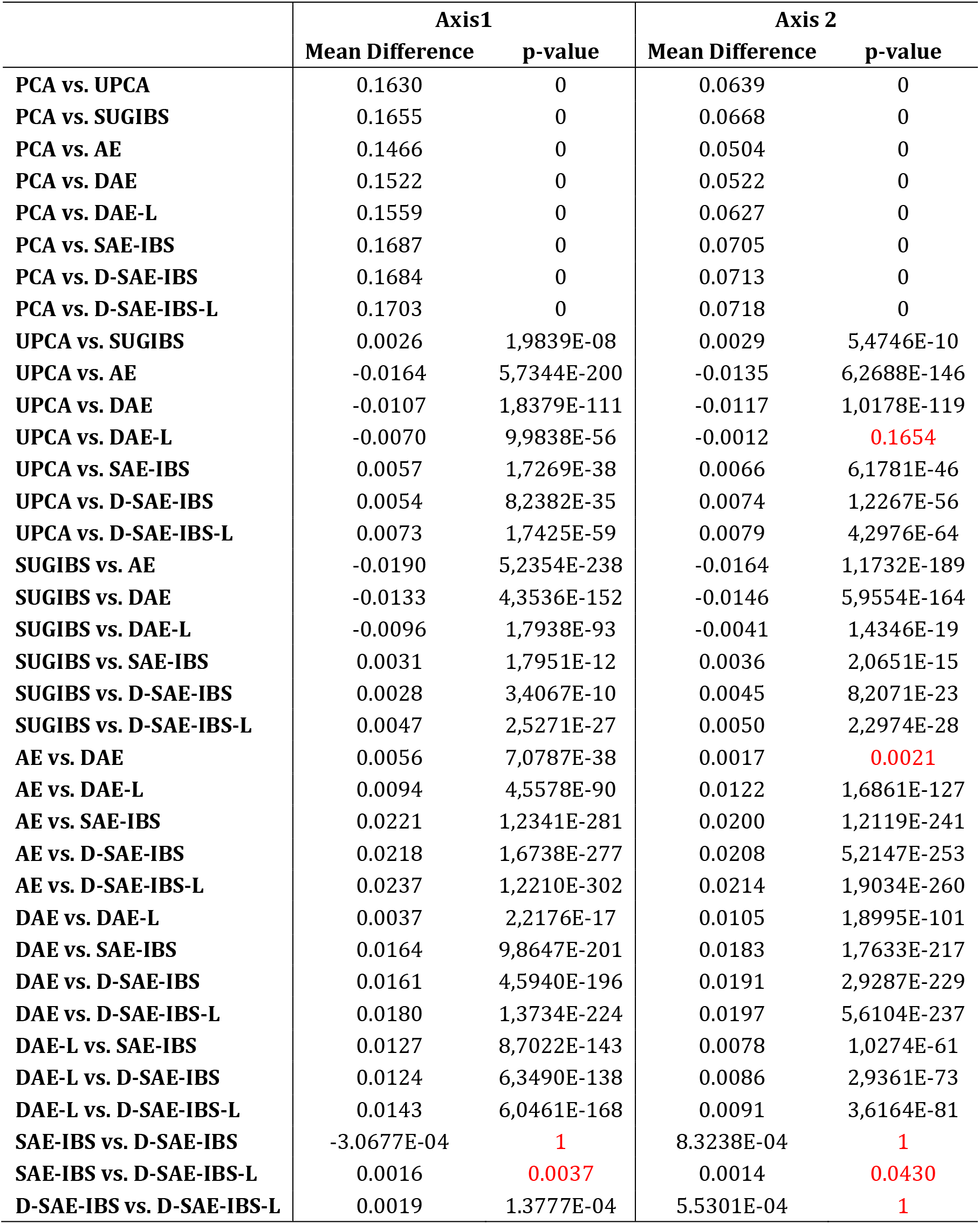
Results of the two-sample t-test on the NRMSD of different methods over 100 simulations for the experiment of missingness. Bonferroni correction method was applied to compute the adjusted significance level, accounting for multiple compassion. The non-significant p-values (p>0.0014) are marked in red.

